# An algorithm for the transformation of the Petri net models of biological signaling networks into influence graphs

**DOI:** 10.1101/2025.01.06.631612

**Authors:** Simon Gamache-Poirier, Alexia Souvane, William Leclerc, Catherine Villeneuve, Simon V. Hardy

## Abstract

A common depiction for biological signaling networks is the influence graph in which the activation and inhibition effects between molecular species are shown with vertices and arcs connecting them. Another formalism for reaction-based models is the Petri nets which has a graphical representation and a mathematical notation that enables structural analysis and quantitative simulation. In this paper, we present an algorithm based on Petri nets topological features for the transformation of the computational model of a biological signaling network into an annotated influence graph. We also show the transformation of the Petri nets model of the beta-adrenergic receptor activating the PKA-MAPK signaling network into its representation as an influence graph.

## 1. Introduction

Petri nets are a formal method from computer science used to model, specify, and study concurrent systems. They serve as both a graphical notation and a mathematical framework. Petri nets are utilized to demonstrate qualitative and quantitative properties of various systems and to simulate their evolution through discrete, continuous, or hybrid processes [1, 2, 3]. This formalism has been applied in diverse fields, including the modeling of biological systems [4, 5, 6]. For instance, Petri nets have been used to demonstrate qualitative properties and reproduce complex hybrid dynamical phenomena in biological systems [7, 8, 9, 10, 11, 12, 13].

In biological modeling, systems are often described using ordinary differential equations (ODE) and analyzed with dynamical systems theory tools [14]. Structural techniques such as pathway analysis and model-checking can complement these analyses [15, 16, 17, 18, 19, 20]. Petri nets combine structure and dynamics, allowing the extraction of reaction network models from differential equations [21]. Using time-course simulation data together with structural analysis of a Petri net biological model, one can construct active state transition diagrams and identify temporal subnets of important dynamical activity [22].

Another common representation in biology is the influence graph [23, 24, 25, 26], which depicts directed interactions between molecular species, labeled with influence types (e.g., activation and inhibition). This representation is useful for illustrating control mechanisms regulating various biological processes in cells, such as metabolic or information flow.

In this paper, we present an algorithm that extracts structural features from a reaction network model specified as a Petri net and creates its influence graph representation. Structural properties such as Petri net invariants, exploration sequences of the bipartite graph, and graph properties like maximal component decomposition are used to extract vertices associated with molecular species and arcs showing influence relationships embedded in the reaction network model. The algorithm is designed to process models of cell signaling systems, where biochemical signals (e.g., hormones, nutrients, endogenous chemicals) initiate production, degradation, release, capture, activation, or inhibition of a network of molecules. These networks mainly consist of proteins such as enzymes (e.g., kinases, phosphatases), second messengers, and other small molecules. Signals travel through the signaling network as enzymes activate or deactivate molecules in cascades until the signal’s target is reached. Signaling networks not only transmit but also process biological signals using regulatory motifs like negative and positive feedback loops, feedforward loops, and more [27, 28, 29]. Understanding how biological signaling is processed by the cell is a key motivation for computational modeling and simulation of these systems [30].

An influence graph, such as the one generated by our algorithm, displays the network and its regulatory motifs, while simulation data demonstrate the biochemical dynamics responsible for signal processing by the network, triggering a cellular response. The influence graph is an unparalleled tool for developing an engineering perspective on biological systems. Figure 1 illustrates the input and output of the transformation algorithm presented in this paper. The input is the Petri net model depicting the regulation of eight executor proteins by cyclin-dependent protein kinase (Cdk) in the eukaryotic cell cycle (Figure 1a, mathematical model in [31]). The algorithm generates an influence graph from this model. Figure 1b displays a section of the resulting influence graph with the influence of Cdk on four executor proteins. This graph reveals four regulatory motifs: two coherent feedforward loops regulating EPP1 and EPP4 and two incoherent feedforward loops regulating EPP2 and EPP3. These motifs enable Cdk to cyclically regulate the synthesis and activation of executor proteins at four different points in the cell cycle. To fully understand cellular control mechanisms: how cells make decisions, how diseases disrupt these processes, and how to intervene pharmacologically, it is essential to grasp the role of individual regulatory motifs in signaling pathways and their interactions [32]. Obtaining this understanding is a central goal of systems biology [33], and influence graphs are a key method to achieve it.

**Figure 1:**
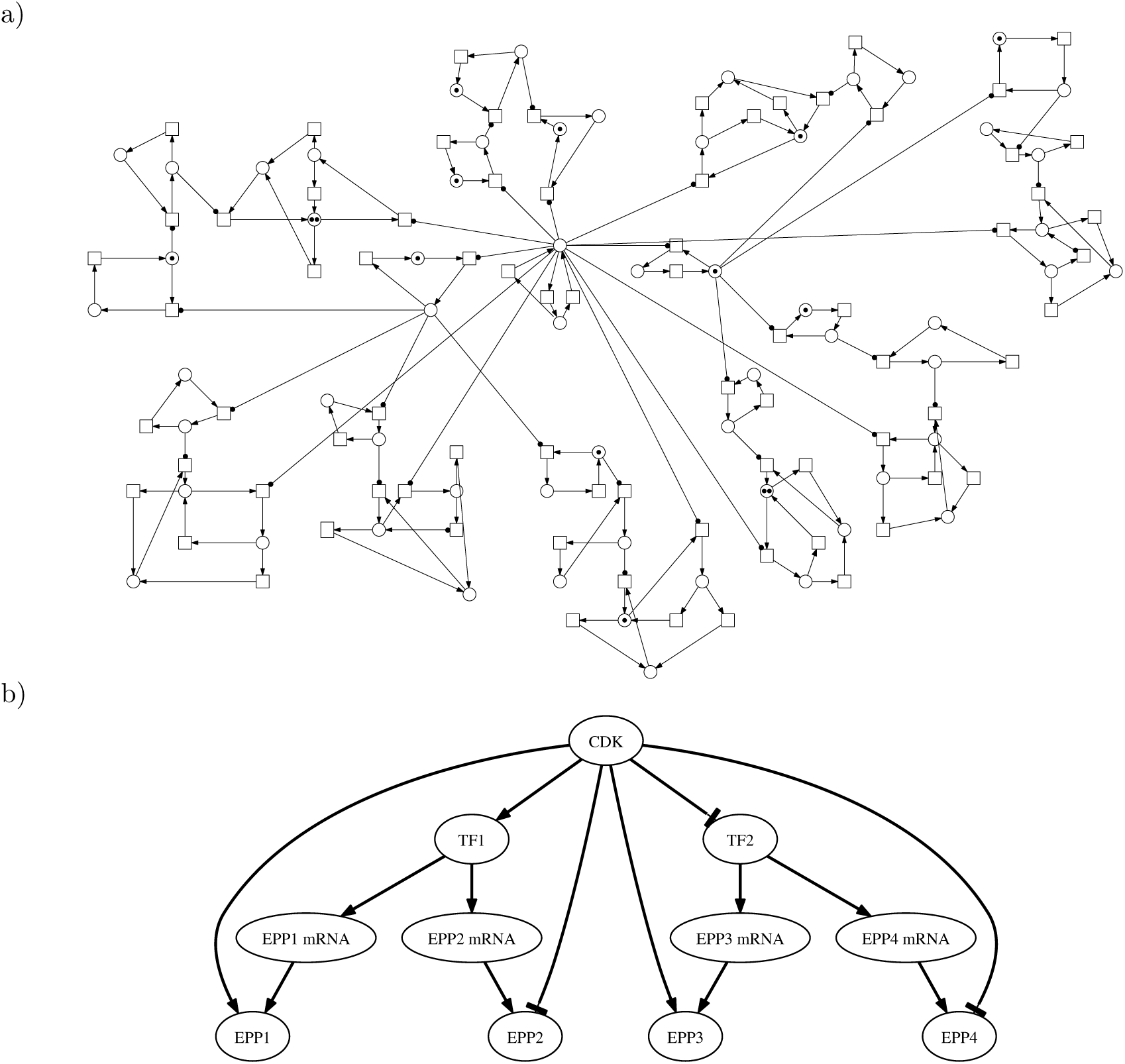
Input and result of the transformation algorithm. In a), the Petri net model of the regulation of eight executor proteins by cyclin-dependent protein kinase (Cdk) in the eukaryotic cell cycle (mathematical model in [31]). In b), part of the influence graph of this model highlighting four feedforward motifs of this network. The full model comprises reactions for the oscillatory regulation of Cdk and four other feedforward motifs.

The transformation algorithm is divided into three phases: preparation, transformation, and annotation. The algorithm detects specific situations and prompts the user with requests for additional information when necessary. This missing information usually corresponds to implicit knowledge in the mathematical model of a biological system. The algorithm can be applied to chemical reaction-based biological models involving binding reactions (where molecules form complexes) or enzymatic reactions (where enzymes transform substrates into products). In differential equations, the kinetic rate of binding reactions is specified by the law of mass action, while the kinetic rate of enzymatic reactions is specified by the Michaelis-Menten formulation if described as a single reaction rather than as three binding reactions.

The first phase of the algorithm performs a preliminary analysis, extracting data from the Petri net model, validating the model if necessary, and conducting a preparatory exploration of the Petri net. The second phase constructs the structure of the influence graph. Finally, the third phase annotates the influence graph with information about active molecular conformations and reactions transmitting the biological signal, determining the type of influence (activation or inhibition). When algorithmically decidable, all transformation and annotation processes are automated. The output of the transformation algorithm can validate the Petri net model and potentially the differential equations model from which it might have originated. In addition, it can be used to create a dynamic representation of the model, allowing visualization of simulation data from a systemic perspective [34, 35, 36].

The paper is organized as follows : Section 2 presents basic terms from Petri net terminology and the marked contextual extension used to model biological systems as reaction network models, along with definitions from graph theory. A detailed introduction to Petri nets can be found in [37]. Section 3 describes the three phases of the proposed algorithm in both textual and algorithmic formats, with simple examples. Section 4 illustrates the proposed algorithm using the Petri net model of the G protein-coupled receptor (GPCR) regulation pathway of the kinases PKA and MAPK (the ODE model was first published in [38]). The final section discusses the different uses of this transformation and potential future improvements of the algorithm.

## 2. Contextual Petri nets terminology and graph theory definitions

In this section, we formally define the marked contextual Petri net and other terms from the Petri net terminology. We continue with other definitions from graph theory that are used in the graph transformation algorithm presented in this paper. A marked contextual Petri net *N*, i.e. a Petri net with read arcs, is a six-tuple, *N* = (*P, T, F, C, W, M*_0_), where *P* = {*p*_1_*, p*_2_*, …, p_n_*} is a finite set of *n* places, *T* = {*t*_1_*, t*_2_*, …, t_m_*} is a finite set of *m* transitions, *F* ⊂ (*P* × *T*) ∪ (*T* × *P*) is a set of arcs, *C* ⊂ *P* × *T* is a set of read arcs, *W* : *F* ∪ *C* → ℕ is a weight function that associates a positive integer to each regular or read arc and *M*_0_ : *P* → ℕ is the initial marking that assigns a non-negative integer to each place. In Petri net theory, the marking represents the number of tokens present in each place. A Petri net is a bipartite graph since *P* ∩ *T* = ∅ and *P* ∪ *T ≠* ∅ (see Definition 2 and Figure 2 for an example).

**Figure 2:**
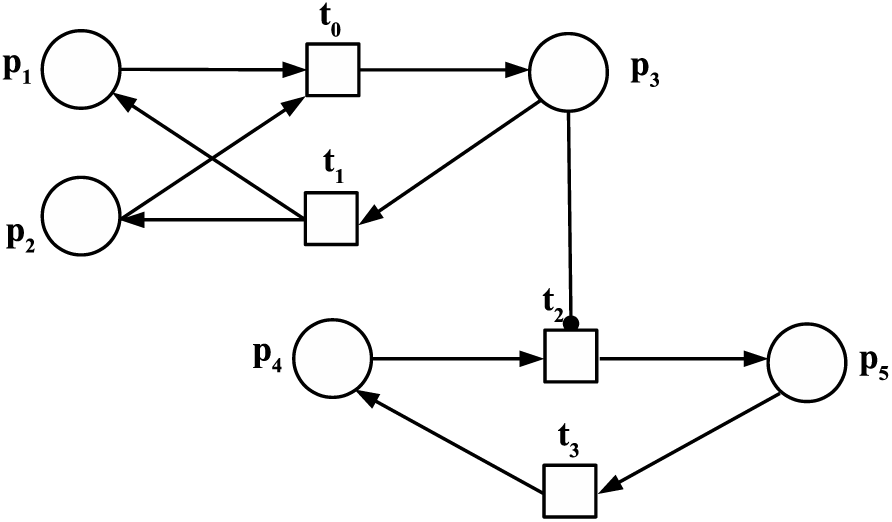
The Petri net *N*_1_ representing a simple network model with four biochemical reactions: the association (*t*_0_)/dissociation (*t*_1_) of two molecular species to form a complex, and the direct enzymatic reaction (*t*_2_) and its reverse (*t*_3_).

For *x* ∈ *P* ∪ *T*, let ^•^*x* := {*y* ∈ *P* ∪ *T* |(*y, x*) ∈ *F* } the preset of *x* (either pre-places or pre-transitions) and *x*^•^ := {*y* ∈ *P* ∪ *T* |(*x, y*) ∈ *F* } the postset of *x* (either post-places or post-transitions). The context of a place *p* is defined as *<u>p</u>* := {*t* ∈ *T* |(*p, t*) ∈ *C*}, and the context of a transition *t* as *t* := {*p* ∈ *P* |(*p, t*) ∈ *C*}. A read arc can only start at a place to connect to a transition.

A function *m* : *P* → ℕ is called the marking of *N*. A transition *t* is enabled at *m* if *m*(*p*) ≥ *w*(*p, t*) for all *p* ∈ *t*∪^•^*t*. Once enabled, *t* can fire, leading to marking *m*^′^, where *m*^′^(*p*) = *m*(*p*)− *w*(*p, t*)+*w*(*t, p*) for all *p* ∈ ^•^*t* ∪ *t*^•^. The firing of a transition can be viewed as the displacement of tokens from input places to output places. By definition, the weight of read arcs is taken into account in the enabling of transitions, but the mark of the places of context *t* of transitions *t* is not modified by the firing of these transitions, hence the term “read arcs”.

The incidence matrix **A** = [*a_ij_*] is a *n* × *m* matrix of integers such that 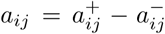, where 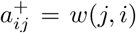 is the weight of the arc from transition *j* to post-place *i* (*j*^•^), and 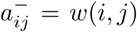 is the weight of the arc from pre-place *i* (^•^*j*) to transition *j*. The incidence matrix excludes the weights of read arcs *C*.

A p- or place invariant (t- or transition invariant) is a *n*-vector *y* (*m*-vector *x*) of integers such that **A***^t^* · *y* = 0 (**A** · *x* = 0). The set of places (transitions) corresponding to non-zero entries in a place invariant *y* ≥ 0 (transition invariant *x* ≥ 0) is termed ‘the support of an invariant’ and is denoted by ∥*y*∥ (∥*x*∥). An invariant *y* is minimal if there is no other invariant *y*_1_ such that *y*_1_(*p*) ≤ *y*(*p*) for all *p*. A t-invariant is called trivial if it is composed of two transitions and their sets of pre- and post-places are symmetrically inverse.

*Example 1:* Figure 2 shows a small Petri net with places *p*_1_ to *p*_5_ and transitions *t*_0_ to *t*_3_ connected by several arcs. The arc between place *p*_3_ and transition *t*_2_ is the only read arc (the head of the arc is a circle instead of an arrow). This Petri net model represents a simple reaction network model. Transitions *t*_0_ and *t*_1_ correspond to a reversible binding reaction, where the molecular species associated with places *p*_1_ and *p*_2_ bind to form the complex *p*_3_ through the firing of *t*_0_ and unbind through the firing of *t*_1_. The read arc indicates that the presence of *p*_3_ is necessary for the reaction *t*_2_ to occur, but that the marking of *p*_3_ is not modified by the firing of the connected transition. This can represent an enzymatic reaction in which *p*_3_ is the enzyme, *p*_4_ is the substrate and *p*_5_ is the product. *t*_3_ is the reverse reaction.

The algorithm also incorporates concepts from graph theory. In general, a graph *G* = (*V, E*) consists of a finite set *V* of vertices and a finite set *E* ⊆ *V* × *V* of edges. The edge (*v*_1_*, v*_2_) ∈ *E* is a link between the vertices *v*_1_ and *v*_2_. The proposed algorithm uses interaction graphs and influence graphs which are variants of general graphs and are defined in subsections 3.2.1 and 3.2.3. The algorithm also performs a complete bipartition decomposition following Definitions 1 to 4.

*Definition 1:* A clique is a subset of vertices in a graph such that every vertex of the clique is connected to one another (i.e. every possible pair of vertices in a clique are adjacent). The subgraph induced by a clique is complete.

*Definition 2:* A bipartite graph *G* = (*V*_1_*, V*_2_*, E*) is a graph whose vertices can be divided into two disjoint and independent sets (for *V*_1_ and *V*_2_, *V*_1_ ∩ *V*_2_ = ∅) such that every edge connects a vertex from one set to a vertex from the other set and never two vertices from the same set. *V*_1_ and *V*_2_ are partitions of *G*.

*Definition 3:* A complete bipartite graph *G* = (*V*_1_*, V*_2_*, E*), or biclique, is a bipartite graph in which every vertex of the first set of vertices is connected to every vertex of the second set. That is for every two vertices *v*_1_ ∈ *V*_1_ and *v*_2_ ∈ *V*_2_, (*v*_1_*, v*_2_) is an edge in *E*.

*Definition 4:* The complete bipartition decomposition of a directed graph *G* is a set of complete bipartite subgraphs *H* = (*V*_1_*, V*_2_*, E*) of *G* such that no 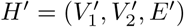 exists such that *V*_1_ is a subset of 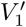 and *V*_2_ is a subset of 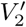. All vertices *v* of *V*_1_ are source vertex in *G* with respect to all target vertex in *V*_2_. Such a subgraph *H* is said to be a maximal complete bipartite component of the graph *G*.

## 3. Description of the proposed algorithm

In this section, we describe the proposed algorithm in detail. Section 3.1 covers the first phase of the algorithm also named the preparatory phase. Section 3.2 describes the second phase of the algorithm, during which the influence graph is created. In this graph, vertices and arcs represent signaling molecules and their signaling interactions, respectively. Finally, Section 3.3 describes the third phase, where the influence graph is annotated. The arcs of the graph are classified as either activation or inhibition, and both vertices and arcs are annotated with elements that allow for the incorporation of the simulation data of a mathematical model to create a dynamic animation of the graph. Table 1 provides an overview of the transformation algorithm highlighting the steps in which the algorithm requests additional information if necessary.

**Table 1:** Overview of the transformation algorithm to create an annotated influence graph from a biological Petri net model.

| Inputs: Petri net model and signaling source place |  |  |
| --- | --- | --- |
| Phase I | Preparation | **1. P-invariant analysis and validation**<br>Sends request to validate similar p-invariants if present<br>Output: Table *placeToInvariant* |
|  |  | **2. Identification of signaling segments**<br>Output: List of signaling segments |
|  |  | **3. Exploration of the Petri net**<br>Output: Table *RFAP* |
| Phase II | Transformation | **4. Building the interaction graph**<br>Uses table *placeToInvariant* to create vertex<br>Uses table *RFAP* to create interaction edges<br>Outputs: $G_{interaction}$ and Table *OE* |
| | | **5. Removal of interaction edges with the signaling segments rule**<br>Iterates through *OE* to remove edges from $G_{interaction}$ with the list of signaling segments<br>Uses the data structure **recorder** that contains several **records** |
| | | **6. Building the influence graph**<br>Decomposes the graph into maximal bipartite components and verifies signal condensation relationships<br>Sends request to choose the interaction partner in molecular complex if undecidable<br>Output: Unannotated $G_{influence}$ |
| Phase III | Annotation | **7. Detecting signaling places and associating them with the vertices**<br>Sends requests for leaves, dissociation states and active states if every place is active or inactive |
|  |  | **8. Associating signaling transitions with influence arcs** |
|  |  | **9. Determining the type of arcs of the influence graph** |
| **Output:** Annotated influence graph |  |  |

### 3.1. Phase I: Preparatory phase and Petri net analysis of the model

In the first phase of the proposed algorithm, known as the preparatory phase, the biological Petri net model is analyzed to identify and validate structural components such as p-invariants and signaling segments. The Petri net model is also explored to determine a sequence of activation steps from a biological perspective, starting from a signaling source to one or more biological targets. The outputs of this phase are several data structures that are used to build the influence graph in the second phase of the algorithm.

#### 3.1.1. P-invariant analysis and validation

The algorithm starts with computing all non-negative, minimal-support p-invariants and t-invariants of the Petri net model (see Section 2) with an implementation of the Fourier-Motzkin (FM) algorithm [39]. The reaction-based biological model of a signaling network usually contains several conservation relationships between state variables (in other words, the sum of these variables is constant). These variables form p-invariants that account for the different biological molecular species that are either transformed variations of the same molecule (e.g., a protein in different phosphorylation states, such as unphosphorylated MAPK, MAPK with a single phosphorylation and MAPK with two phosphorylations) or molecules that are part of several molecular complexes (e.g., the Ca^2+^ ion in complex with different proteins, such as CaM.Ca_4_, PDE.Ca and AC.Ca) [40]. In these two cases, the sum of the concentrations is constant, indicating a conservation relationship. The algorithm presented in this paper relies on this relationship between p-invariants and biological entities to identify the vertices of the influence graph. In Example 2 and Figure 3, we show an example of the result of p-invariant and t-invariant analysis on the Petri net *N*_1_.

**Figure 3:**
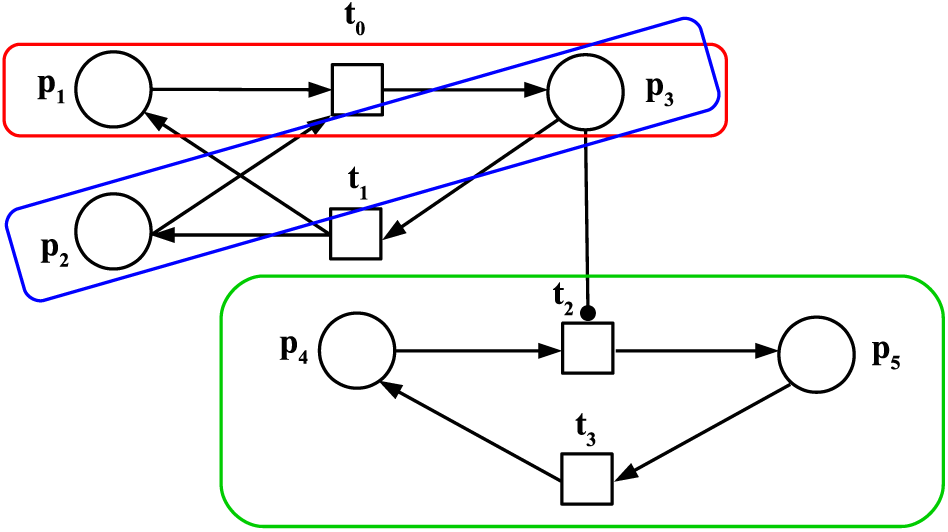
Result of the p-invariant analysis performed on the Petri net *N*_1_. In red, blue and green are the three minimal p-invariants *y*_1_, *y*_2_ and *y*_3_ respectively.

*Example 2:* Consider the Petri net *N*_1_ in Figure 3 and its incidence matrix **A**. By solving the equation **A***^t^* ·*y* = 0, we identify three minimal p-invariants: ∥*y*_1_∥ = {*p*_1_*, p*_3_}, ∥*y*_2_∥ = {*p*_2_*, p*_3_} and ∥*y*_3_∥ = {*p*_4_*, p*_5_}. Each p-invariant corresponds to a conservation relationship between molecular species. Solving the equation **A** · *x* = 0, we find the following t-invariants: ∥*x*_1_∥ = {*t*_0_*, t*_1_}, ∥*x*_2_∥ = {*t*_2_*, t*_3_}. These two t-invariants are trivial. In biological terms, the molecular species represented by *y*_1_ and *y*_2_ interact to form a complex, which acts as an active enzyme. This enzyme modifies the molecular species *y*_3_, transforming it from state *p*_4_ to state *p*_5_.

*Definition 5: E_p_* denotes the set of all p-invariants place *p* belongs to.

After the p-invariant analysis is performed, an entry is made in the lookup table *placeT oInvariant* for each place, listing the p-invariants this place belongs to.

*Definition 6: placeT oInvariant* denotes a lookup table such that *placeT oInvariant*(*key*) = *E_p_*. The value associated with each key *p* is the set of p-invariants *E_p_* to which the place *p* belongs: *placeT oInvariant*(*p*) = *E_p_*. An example of the lookup table *placeT oInvariant* for the previous p-invariant analysis (Figure 3) is given in Table 2.

**Table 2:** The lookup table *placeT oInvariant* for the Petri net of Figure 3.

| Place | $E_p$ |
| --- | --- |
| $p_1$ | $y_1$ |
| $p_2$ | $y_2$ |
| $p_3$ | $y_1, y_2$ |
| $p_4$ | $y_3$ |
| $p_5$ | $y_3$ |

In Heiner et al. [40], the authors suggested that every p-invariant corresponds to a valid biological component of a model. However, we found a counter-example to this statement, presented in Figure 4. This Petri net model includes 4 bidirectional biochemical reactions involving molecular species *A* (an inhibitor), *B* (a second inhibitor) and *C* (an enzyme). Initially, *B* is in complex with two molecules of *C*, inhibiting them. *B* also has two binding sites for *A*. When the complex *BC*_2_ binds to two molecules of *A*, the inhibitor *B* is repressed, and the enzyme *C* is released in its active form. One would expect the structural analysis of this model to identify three p-invariants corresponding to the molecular species *A*, *B* and *C*. However, this model has four p-invariants ({*BC*_2_, *BC*_2__*A*, *BC*_2__*A*_2_, *BC*_*A*_2_, *B*_*A*_2_}, {*BC*_2_, *BC*_2__*A*, *BC*_2__*A*_2_, *BC*_*A*_2_, *C*}, {*A*, *BC*_2__*A*, *BC*_2__*A*_2_, *BC*_*A*_2_, *B*_*A*_2_} and {*A*, *BC*_2__*A*, *BC*_2__*A*_2_, *BC*_*A*_2_, *C*}). Despite the names of the places, it is mathematically impossible to determine from an invariant analysis alone whether molecule *A* is bound to *B*, *C*, or both. The structural analysis identifies every possibility. However, to obtain the correct vertices for the biological influence graph from a biological Petri net model, only p-invariants corresponding to valid biological entities must be considered. Thus the results from the p-invariant analysis have to be validated. We found that the support of an invalid p-invariant shares a high degree of similarity with the support of a valid p-invariant. To assist in detecting invalid p-invariants, the algorithm detects similar p-invariants using an arbitrary threshold of 60%.

**Figure 4:**
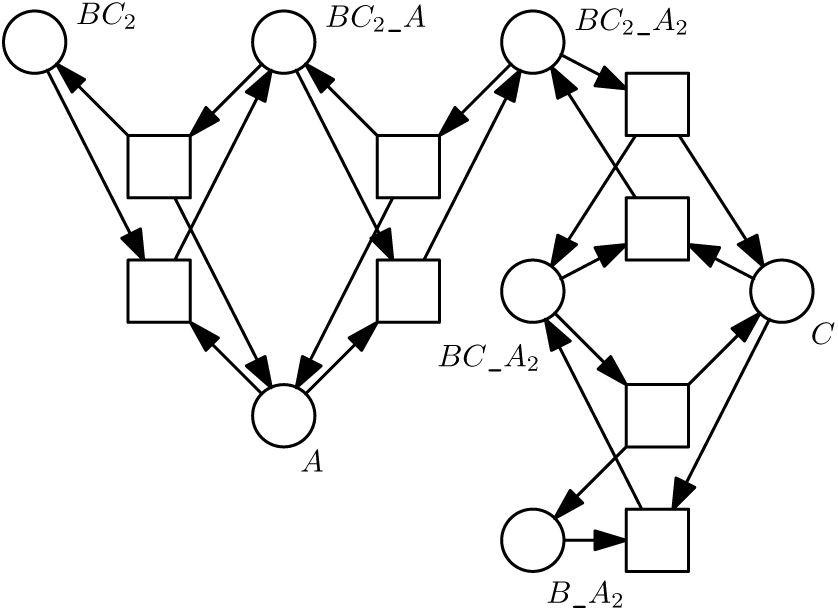
Petri net model of biochemical reactions with more p-invariants (4) than real molecular species (3: A, B and C). This example justifies the need for a validation of the detected p-invariants.

*Definition 7:* Two p-invariants *y*_1_ and *y*_2_ are considered similar if |(∥*y*_1_∥ ∩ ∥*y*_2_∥)|*/max*(|∥*y*_1_∥|, |∥*y*_2_∥|) *>* 60%.

When two similar p-invariants are detected, a request is sent to a user to determine if one of the p-invariants is invalid. The p-invariant selected as invalid is then removed from table *placeToInvariant*.

#### 3.1.2. Identification of signaling segments

In the second phase of the algorithm, an interaction graph is built to track the molecular signal as it is transduced through the signaling network, starting from a place acting as a signaling source. To prevent the signal from traveling backwards, a transformation rule defined in subsection 3.2.2 removes edges from the initial interaction graph built by the previous rule based on specific sets of transitions, which we call signaling segments. These segments consists of sets of transitions that operate as units connected together to transmit biological signals throughout the network. In this rule of the first phase of the algorithm, signaling segments are identified from the t-invariants of subnets of *N*. These subnets comprise the places of a p-invariant and the transitions connecting them in the Petri net model. Consequently, there are as many subnets in the Petri net model as there are p-invariants.

*Definition 8:* Let 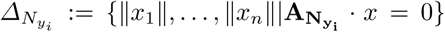 be the set of the *n* minimal t-invariant support sets of the subnet *N_yi_*. The set of signaling segments of subnet *N_yi_* is defined as 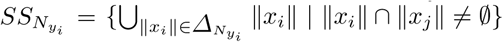. A signaling segment is a set of transitions created by the union of every minimal t-invariant supports of the subnet having at least one transition in common. A p-invariant subnet can have more than one signaling segment.

The results of this rule are stored in the sets 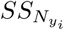, one for each p-invariant, which will be used in the second part of the algorithm.

*Example 3:* In the Petri net *N*_1_, the three subnets *N_y_*_1_, *N_y_*_2_ and N*_y_*_3_ are analyzed, one for each p-invariant. For example, the subnet *N_y_*_1_ for the p-invariant *y*_1_ contains the places *p*_1_ and *p*_3_ connected to the transitions *t*_0_ and *t*_1_. These transitions form the support of the only t-invariant of the subnet *N_y_*_1_, and this set is a signaling segment. The sets of signaling segments of this model are 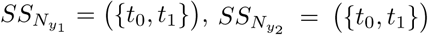 and 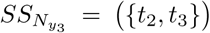. Due to the simplicity of this example, the signaling segments correspond to trivial t-invariants. However, in more complex nets like the model *N*_2_ of Subsection 3.2.3 and the model *N_GP CR_* from Section 4, the set of a signaling segment might contain several trivial t-invariants merged together if they have transitions in common.

#### 3.1.3. Exploration of the Petri net model starting from the signaling source of the network to create an exploration sequence

The final step of the preparatory analysis in the first phase of the algorithm involves an exploration sequence of the Petri net, moving from place to place through transitions. Starting from a signaling source place provided by the user, the algorithm performs a depth-first search (DFS) of the Petri net. This sequence is recorded in another data structure: a lookup table named *RFAP* (for ReachedFromAllPlaces) is built as the Petri net is explored and places are visited. The exploration sequence will be used in the transformation phase to build the interaction graph. From the visited place *p_source_*, every post-transition *t* ∈ *p_source_*^•^ is visited, and their post-places *p_target_* ∈ *t*^•^ are reached. To the three vertices *p_source_*, *p_target_* and *t*, a number corresponding to the *step* of exploration is added, forming a quadruplet. The exploration then continues from these newly visited places. Each place can be visited only once.

*Definition 9:* A quadruplet *q_i_* is composed of a source place, a target place, a transition and a step. It is stored in a list as [*p_source_, p_target_, t, step*]. The step is a number referring to the step rank in the depth-first search. An example of a quadruplet of Petri net vertices is shown in Figure 5.

**Figure 5:**
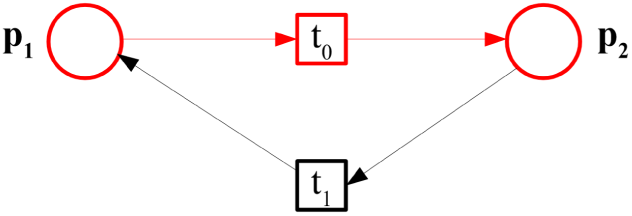
Example of the quadruplet [*p*_1_*, p*_2_*, t*_0_, 0] (in red). In this case, the exploration step is 0 because it is the first step from the specified signaling source *p*_1_.

An entry is made in the lookup table *RFAP* for each place *p*. The entries in the lookup table *RFAP* are a list of the quadruplets in which *p* is the source place.

*Definition 10: RFAP* denotes a lookup table such that *RFAP* (*key*) = set of quadruplets. The value associated with each key *p* is a set of quadruplets whose source place is *p*.

*Example 4:* Let *p*_1_ be the place of the signaling source of the model. This is the place where the exploration of the Petri net *N*_1_ starts. Table 3 shows the *RFAP* lookup table for the DFS of the Petri net *N*_1_. A complete description of how the table is built is provided in the supplementary material.

**Table 3:**
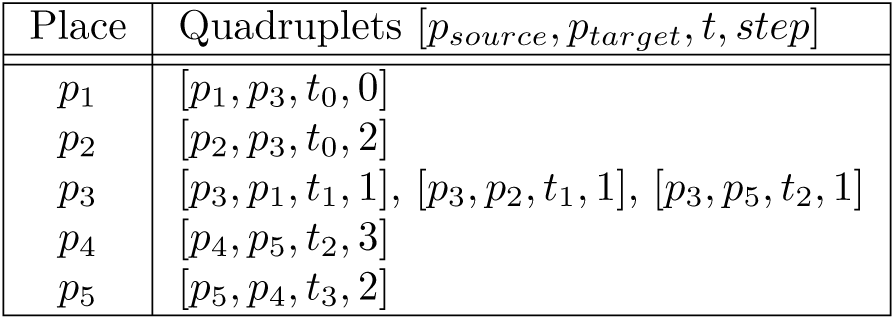
Places and their quadruplets in the lookup table *RF AP* for the Petri net *N*_1_.

| Place | Quadruplets $[p_{source}, p_{target}, t, step]$ |
| --- | --- |
| $p_1$ | $[p_1, p_3, t_0, 0]$ |
| $p_2$ | $[p_2, p_3, t_0, 2]$ |
| $p_3$ | $[p_3, p_1, t_1, 1], [p_3, p_2, t_1, 1], [p_3, p_5, t_2, 1]$ |
| $p_4$ | $[p_4, p_5, t_2, 3]$ |
| $p_5$ | $[p_5, p_4, t_3, 2]$ |

The first phase of the proposed algorithm concludes with the exploration of the Petri net model. During this analysis, similar p-invariants are detected to verify their biologically validity. Additionally, three different data structures are created from the Petri net of the biological signaling model: 1) the lookup table *placeT oInvariant* containing p-invariant information, 2) a list of signaling segments of the Petri net and 3) the *RFAP* lookup table with an exploration sequence of the places and transitions of the Petri net from a designated source place. This information is used in the second phase of the algorithm to build the influence graph. The preparatory phase of the proposed algorithm is summarily described in Algorithm 1. The algorithm is also provided in more formal detail in Algorithm S1.

##### Algorithm 1 Preparatory phase and Petri net analysis (phase I)

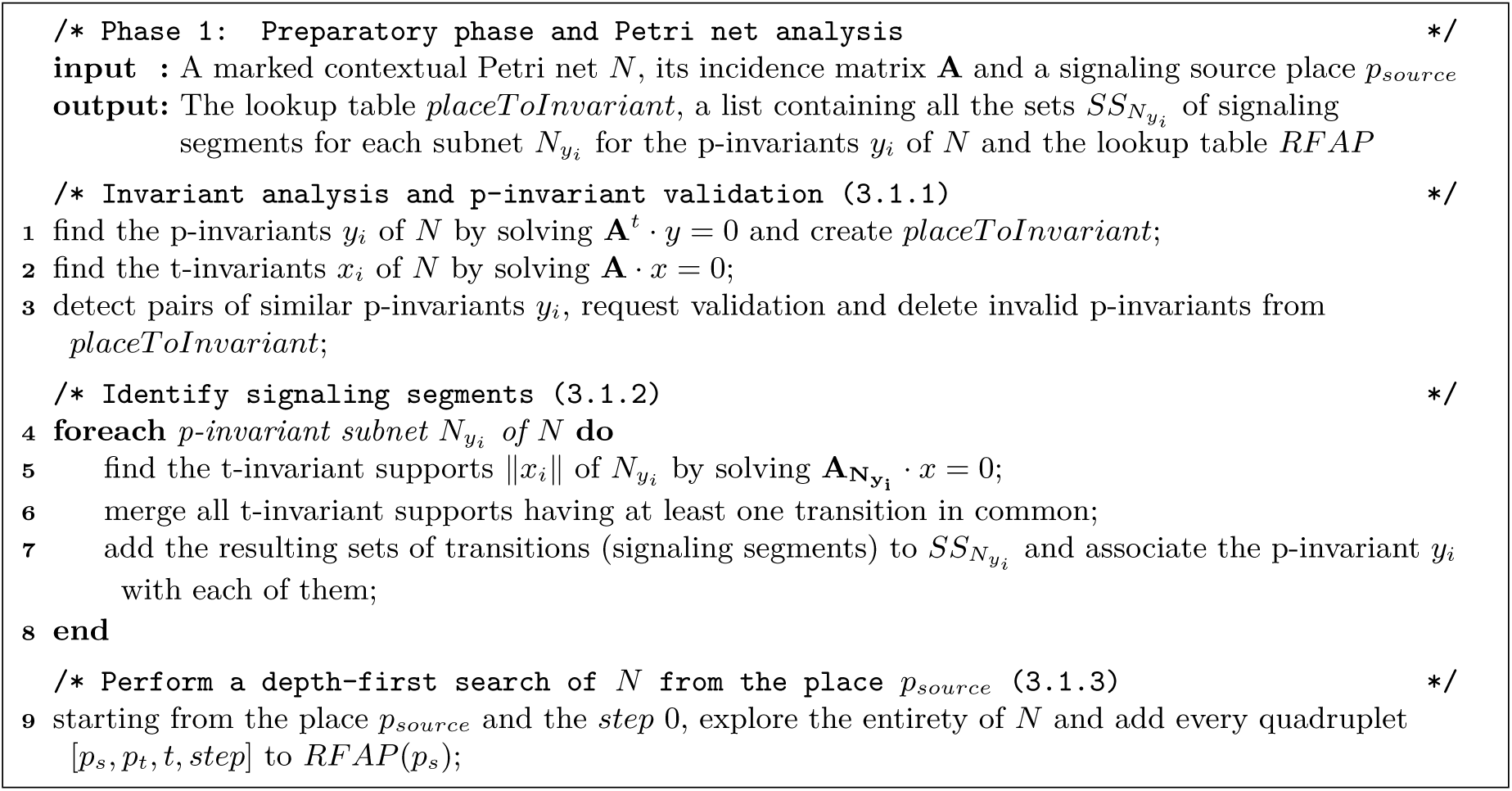

### 3.2. Phase II: Transforming the Petri net model into an influence graph

In the second phase of the Petri net transformation algorithm, an influence graph is constructed from the biological Petri net model through three transformation rules and using the three data structures created in the previous phase. In the first rule of the second phase, the intermediate representation of the interaction graph is built, establishing the relationships between the p-invariant sets of the Petri net model. After being created, some edges in the interaction graph are removed following the second rule based on signaling segments. The third and final rule transforms the interaction graph into the influence graph. This graph will be annotated in the third phase of the algorithm.

#### 3.2.1. Building the interaction graph of the Petri net model

The first rule of the second phase of the algorithm is to transform the Petri net model into an interaction graph. An interaction graph is a directed multigraph in which no self-loops are permitted, and multiple edges between any two vertices are possible. The interaction graph *G_interaction_* = (*V, E*) consists of a finite set *V* of vertices and a finite set *E* ⊆ *V* × *V* of edges. The edge (*v*_1_*, v*_2_) ∈ *E* is a link between the vertices *v*_1_ and *v*_2_. In the interaction graph, a vertex is associated with one or several p-invariants, and an edge is associated with a unique quadruplet (see definition 9). If we consider *v_i_* a vertex of *G_interaction_*, we denote *Inv*-*G_interaction_*(*v_i_*) as the unique set of p-invariants associated with *v_i_*. If we consider *e_i_* an edge of *G_interaction_*, we denote *Tr*-*G_interaction_*(*e_i_*) as the quadruplet associated with *e_i_*.

*Definition 11:* For each different p-invariant set *E_p_* in the lookup table *placeT oInvariant*, we build a vertex *v_i_* in the interaction graph *G_interaction_* and associate *E_p_*(set of p-invariants) with the created node such that *Inv*-*G_interaction_*(*v_i_*) = *E_p_*.

*Definition 12:* The support of an interaction vertex *v_i_* is defined as the set of all the places *p* such that *C*[*p*] → *Inv*-*G_interaction_*(*v_i_*). In other words, the support of *v_i_* is the set containing all the places associated with the value *E_p_* in table *placeT oInvariant*. This value is also equal to *Inv*-*G_interaction_*(*v_i_*).

We denote ||*v_i_*|| as the support of the vertex *v_i_*. Unlike the support of a p-invariant in a Petri net, a place *p* can be present in the support of only one vertex of the interaction graph.

*Example 5:* For the Petri net *N*_1_, vertices are created for the four distinct values *E_p_* in the lookup table *placeT oInvariant* (Table 2). Table 4 shows the list of vertices *v_i_* from the interaction graph representation of *N*_1_ with their set *Inv*-*G_interaction_*(*v_i_*) and their associated support.

**Table 4:**
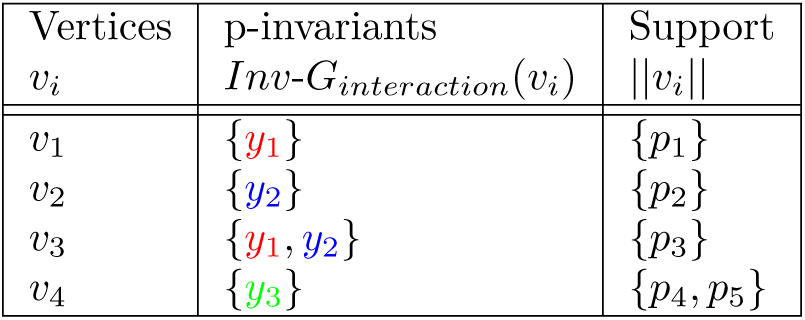
Interaction graph vertices, their associated p-invariant set and their support of the interaction graph created from the Petri net *N*_1_.

| Vertices<br>$v_i$ | p-invariants<br>$Inv-G_{interaction}(v_i)$ | Support<br>$\vert \vert v_i\vert \vert $ |
| --- | --- | --- |
| $v_1$ | $\{y_1\}$ | $\{p_1\}$ |
| $v_2$ | $\{y_2\}$ | $\{p_2\}$ |
| $v_3$ | $\{y_1, y_2\}$ | $\{p_3\}$ |
| $v_4$ | $\{y_3\}$ | $\{p_4, p_5\}$ |

After the creation of the vertices, the edges of the interaction graph are created. For the creation of the edges, each vertex is iteratively considered as a possible source vertex. The places in the support of the considered vertex are used as keys to access quadruplets in the *RFAP* lookup table. Each quadruplet is then assessed: from the target place of a quadruplet, which is the second element of the quadruplet, the associated vertex is identified. This becomes the target vertex of the new edge. Unless the source vertex and the target vertex are the same (no loops are allowed in the interaction graph), an edge is created between these two vertices, and the quadruplet is associated with it. When an edge is added to the interaction graph, an entry is made in another lookup table identified as *OE* (for OrderedEdge). Each entry in the lookup table *OE* contains the step number (the fourth entry of the quadruplet) and the set of quadruplets created at this step during the exploration of the Petri net. The set of quadruplets for each step is *Q_step_*.

*Definition 13:* An edge *e* exists from vertex *v_a_* to vertex *v_b_* in the interaction graph *G_interaction_* if there is a quadruplet *q* = [*p_source_, p_target_, t, step*] in *RFAP* such that *p_source_* is contained in ||*v_a_*|| and *p_target_* is contained in ||*v_b_*||. We associate *q* to the interaction graph edge *e* (*Tr*-*G_interaction_*(*e*) = *q*).

*Definition 14: OE* denotes a lookup table such that *OE*(*key*) = *Q_step_*. The value associated with each key *step* is a set of quadruplets created at this step.

*Example 6:* Five edges are added to the interaction graph of the Petri net *N*_1_. They are illustrated in Figure 6. Each edge is associated with a quadruplet from Table 3. The source and target vertices of an edge are determined from the source and target places of the quadruplet and the corresponding support of the vertices. Because the source and target places of the quadruplets [*p*_4_*, p*_5_*, t*_2_, 3] and [*p*_5_*, p*_4_*, t*_3_, 2] are related to a single interaction vertex (*v*_4_), they are not converted into edges in the interaction graph. Table 5 presents an example of the lookup table *OE* with the remaining five quadruplets for the Petri net *N*_1_.

**Figure 6:**
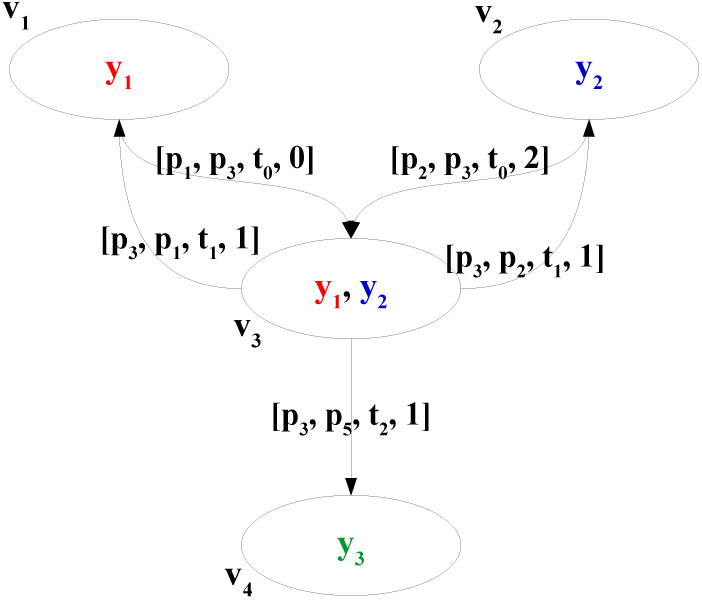
Interaction graph built from the Petri net *N*_1_ after the execution of the first rule of the second phase with its vertices and their associated p-invariants, and its edges and their associated quadruplets.

**Table 5:** The lookup table *OE* for the running Petri net *N*_1_, representing each step and the quadruplets of the associated interaction graph edges.

| Step | Quadruplets of the interaction graph edges |
| --- | --- |
| 0 | $[p_1, p_3, t_0, 0]$ |
| 1 | $[p_3, p_1, t_1, 1], [p_3, p_2, t_1, 1], [p_3, p_5, t_2, 1]$ |
| 2 | $[p_2, p_3, t_0, 2]$ |

#### 3.2.2. Removal of edges from the interaction graph with the signaling segments rule

The final objective of the transformation algorithm is to represent signaling pathways as an influence graph showing the path of the signal. In these cellular systems, most chemical reactions come with their reverse counterpart (e.g. activation and inactivation through phosphorylation and dephosphorylation). As the molecular signal travels from a source through different signaling pathways to one or more molecular targets, the initial perturbation is thought to convey the signal further downstream. When the reverse reactions occur, this likely corresponds to the system trying to return to its initial equilibrium (after activation often comes inactivation). To transform the interaction graph into an influence graph, edges corresponding to these reverse reactions that do not transmit the signal are identified and removed. In the algorithm, this pruning of the graph is done according to the signaling segments rule, using a history of which signaling segments are transducing the signal, from and to which p-invariants and at which step. Three data structures are put in place to track this signal transduction history. The first of these data structures is the record.

*Definition 15:* A record denotes a list of sets containing p-invariants. For each step of the exploration of the Petri net model, there is a set of p-invariants in the record. A record is used to store the sets of p-invariants that are reached at certain steps. The record is initialized with empty sets.

*Example 7:* For a Petri net model whose exploration takes *n* steps, the initial record is as follows: record = [{p-invariants set at step 0}, {p-invariants set at step 1}, {p-invariants set at step 2}, …, {p-invariants sets at step n}] = [{}_0_, {}_1_, {}_2_, …, {}*_n_*]

The record data structure is used repeatedly in this rule of the algorithm. A `record` is associated with one signaling segment (see Definition 8) to form a pair that we identify as a `recorder`.

*Definition 16:* A `recorder` denotes a data structure pairing the signaling segment of a p-invariant subnetwork with a `record`.

*Example 8:* `recorder` = [signaling segment, `record`]

A final data structure is necessary for the execution of the signaling segment rule: the lookup table `invariantToRecorders` in which each recorder is associated with a p-invariant of the Petri net model. One p-invariant can be associated with multiple recorders.

*Definition 17:* The table `invariantToRecorders` is a multiple-value lookup table such that `invariantToRecorders`(key) = list of recorders. The value associated with each key p-invariant is a list of recorders.

*Example 9:* `invariantToRecorders` = ((p-invariant_1_, (recorder_1_, recorder_2_)), (p-invariant_2_, (recorder_3_)), …)

The three data structures – `record`, `recorder` and `invariantToRecorders` – are used only to process the interaction graph within this rule and will not be used in later steps of the algorithm. In the Petri net *N*_1_, these data structures are initialized as follows. There are three p-invariants *y*_1_, *y*_2_ and *y*_3_ in this model, thus there are three entries in the table `invariantToRecorders`. Since the sets of signaling segments of these p-invariants subnets 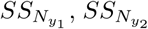 and 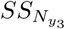 contain only one signaling segment each, there is only one `recorder` per p-invariant in the table. Finally, the record of each recorder is initialized with three empty sets since this Petri net was explored in three steps. Some of these records will be filled as the rule is applied and edges of the interaction graph are removed. We now explain this rule, first in generic terms and then through its application to the Petri net *N*_1_ in Example 10.

When applying the signaling segments rule to an interaction graph, the graph edges are considered based on the exploration step. Thus, the edges at step 0 are first considered, then the edges at step 1, and so on. When an edge is evaluated, the p-invariants from the source place and from the target place of that edge are identified using the function `implicitConnectedInv` as defined in Algorithm 2.1. This function handles vertices associated with more than one p-invariant, which typically involves molecular complexes. These source and target p-invariants are used to access lists of recorders stored in the lookup table `invariantToRecorders`. The correct recorders within these lists are selected by verifying if the transition associated with the edge is present in the signaling segment of the recorder (see function `getRecorder` in Algorithm 2.1). The considered edge will be removed from the interaction graph if one of the source p-invariants is already present in the record of the target p-invariants. If this condition is met, all edges sharing the same source place and transition are also removed from the interaction graph. This indicates that the signal has already traveled along this signaling segment in one direction at a previous step, hence the edges associated with the reverse reactions are removed. Otherwise, the edge is kept, and the target p-invariants are added to the record of the recorder of the source p-invariants. In this manner, the recorders of the model p-invariants are gradually filled and used to test the eligibility of the edges at the remaining steps.

Iterating through the edges of the graph based on the exploration step is crucial in applying this rule. If the signal travels again on a signaling segment that was taken at an earlier step, the corresponding edge, along with any other edge having the same source place and transition (at the same step), is removed from the interaction graph. This removal indicates that these interactions do not transduce the signal, but rather correspond to the system returning to its equilibrium. It is also important to note that this rule does not remove the longer loops created by edges that do not share the same signaling segment. These loops, like negative feedback mechanisms, are important for biological regulation and are thus preserved in the interaction graph.

*Example 10:* Let’s apply the signaling segments rule to the Petri net *N*_1_ and possibly remove certain edges of the interaction graph. The process is formally detailed in Algorithm 2.1. The resulting interaction graph is shown in Figure 7. The state of the lookup table `invariantToRecorders` after the application of the rule is shown in Table 6. A complete description of this process is available in the supplementary material.

**Figure 7:**
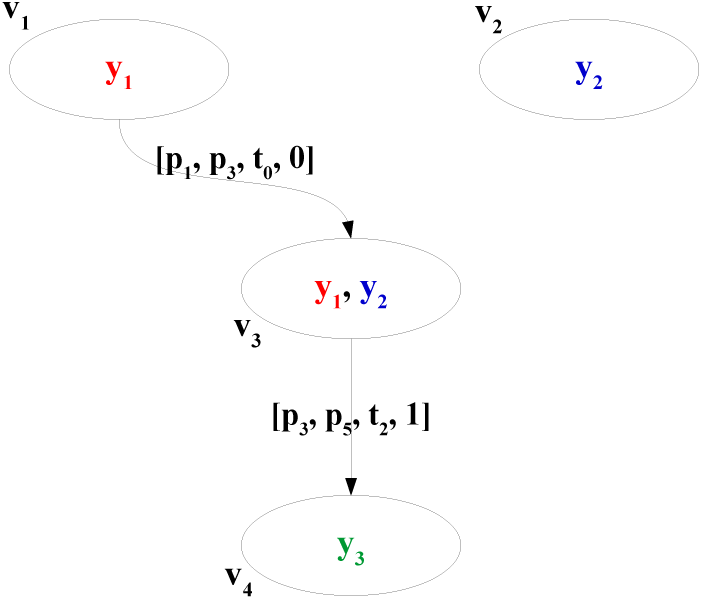
Interaction graph of the Petri net *N*_1_ with the remaining edges and their associated quadruplet after the application of the signaling segments rule when the place *p*_1_ is considered the signaling source of the model.

**Table 6:** The lookup table `invariantToRecorders` for the Petri net *N*_1_ after applying the signaling segments rule. The p-invariant *y*_2_ is added to the `record` of the signaling segment {*t*_0_, *t*_1_} of the p-invariant *y*_1_ when the edge from *v*_1_ to *v*_3_ is considered. This means that the signal travels from *y*_1_ to *y*_2_ using this signaling segment. When edges at later steps between *v*_1_ and *v*_3_ and *v*_2_ and *v*_3_ are considered, they are removed from the interaction graph because the quadruplet transitions are in the same signaling segment.

| P-invariant | Recorder |  |
| --- | --- | --- |
|  | Signaling segment | Record |
| $y_1$ | $\{t_0, t_1\}$ | $[\{y_2\}, \{\}, \{\}]$ |
| $y_2$ | $\{t_0, t_1\}$ | $[\{\}, \{\}, \{\}]$ |
| $y_3$ | $\{t_2, t_3\}$ | $[\{\}, \{\}, \{\}]$ |

#### 3.2.3. Building the influence graph from the interaction graph

The final rule of the second phase of the algorithm transforms the interaction graph into an unannotated version of the influence graph. At this stage of the algorithm, influences of molecular species on others are inferred from molecular interactions. This rule processes molecular complexes to distinguish between direct and indirect interactions. For example, if a complex of three molecules is assembled through binding reactions in the model (e.g. a molecular complex formed of the p-invariants *y_A_*, *y_B_* and *y_C_*), a subset or all the possible combinations formed by the three p-invariants are present in the interaction graph (the vertices *y_A_y_B_*, *y_A_y_C_*, *y_B_y_C_* and *y_A_y_B_y_C_*). This could lead to the creation of influence edges in one direction or the other between the three pairs of p-invariants (*y_A_* → *y_B_*, *y_A_* → *y_C_* and *y_B_* → *y_C_*, and their reverse). However, a direct molecular interaction between all three pairs of molecules forming the complex through binding domains is unlikely. In this situation, it is more probable that only one molecule can bind to the two other partners through distinct binding sites. These other two partners do not bind together on their own and thus should not have a direct influence on one another in the influence graph. In other words, their interaction is indirect. To distinguish between indirect interactions in a molecular complex and direct interactions through binding sites, this rule uses graph properties. In the end, direct interactions are preserved while redundant or indirect interactions are removed from the influence graph. As a result of this processing, the vertices of the influence graph have only one p-invariant associated with them, and arcs have a set of quadruplets. This influence graph will be annotated in the third phase of the algorithm.

An influence graph *G_influence_* = (*V, E*) consists of a finite set *V* of vertices and a finite set *E* ⊆ *V* ×*V* of arcs. The arc (*v*_1_*, v*_2_) ∈ *E* is a directed arc from vertex *v*_1_ to vertex *v*_2_, and means that *v*_1_ has some influence over *v*_2_. There can be only one arc between two vertices per direction. If we consider *v* a vertex of the influence graph, we denote *Inv*-*G_influence_*(*v*) the only p-invariant associated with *v* and ||*v*|| the support of this p-invariant. If we consider *e* an arc of the influence graph, we denote *Tr*-*G_influence_*(*e*) the set of quadruplets associated with *e*. Each arc also has a type of influence. *I* : *E* → *IT* is a type function that associates an influence type to an arc, where *IT* is a finite set of influence types, *IT* = {*it*_1_*, it*_2_*, …, it_m_*}. The influence type of the edges of the influence graph will be assigned by a later transformation rule (subsection 3.3.3). In the interaction graph, a vertex can be associated with more than one p-invariant (the cardinality of the set *Inv*-*G_interaction_*(*v_i_*) for any *v_i_* is greater than or equal to one). However, a vertex in the influence graph is associated with only one p-invariant (the cardinality of the set *Inv*-*G_influence_*(*v_j_*) for any *v_j_* is equal to one). To create the influence graph, the edges from the interaction graph are processed. Since these edges can connect vertices associated with multiple p-invariants, this rule extracts the corresponding influence arcs between single p-invariants. We call the central concept of this transformation rule signal condensation. This is the reduction of interactions into direct molecular influences. Because only binding reactions can create complexes with multiple p-invariants, the condensation of signals is only needed for edges that are associated with binding reactions and not enzymatic reactions.

The transformation rule that processes the interaction graph to create the influence graph proceeds one exploration step at a time, executing five main processing steps as one iteration for each exploration step of the model starting at the step 0. The **first processing step** is to group interaction vertices and edges by the current exploration step to create an interaction subgraph.

The **second processing step** is to condense the signal in each vertex of the exploration step interaction subgraph with the previously identified signal condensation relationships (see the fourth step). There is no signal condensation relationship at the first iteration. Condensing the signal in each vertex of the subgraph is done by removing p-invariants that are codomain with another p-invariant (e.g. if in a condensation relationship, *y*_1_ is the codomain of *y*_2_, then in each vertex where *y*_1_ and *y*_2_ are present, remove *y*_1_). The information is stored in a data structure as signal condensation relationships are detected following the conditions in Definition 18.

*Definition 18:* The signal from the vertex *v*_1_ to the vertex *v*_2_ (associated with p-invariants *y*_1_ and *y*_2_ respectively in the influence graph) is said to be condensed by *v*_2_ under two conditions. First, the edge must not be associated with an enzymatic reaction. The second condition is if there is an edge between the two vertices in the interaction graph that are each associated with sets containing the p-invariants *y*_1_ and *y*_2_ where either source or target vertex contains only one of these p-invariants (the three possible combinations for *Inv*-*G_interaction_*(*v*_1_) and *Inv*-*G_interaction_*(*v*_2_) are {*y*_1_} and {*y*_2_}, {*y*_1_} and {*y*_1_*, y*_2_}, and {*y*_1_*, y*_2_} and {*y*_2_}). A p-invariant that is the sole p-invariant of a vertex is called a core p-invariant. In the resulting condensation relationship from *v*_1_ to *v*_2_, *y*_1_ becomes a codomain of *y*_2_ and *y*_2_ becomes a domain of *y*_1_.

The **third processing step** extracts influence arcs from the interaction graph edges. It starts by decomposing the exploration step interaction subgraph into a set of its maximal bipartite components using the Bron-Kerbosch complete bipartition decomposition algorithm [41]. A bipartite component is a complete bipartite graph (Definition 3) and the set of all maximal complete bipartite components respects Definition 4. Each component is a subgraph with two sets of vertices: sources and targets.

Following the complete bipartition decomposition of the exploration step subgraph, the listed maximal components are analyzed one by one by going through the operations (a) to (e) to extract new influence arcs.

(a) Removing p-invariants from vertices at the second step because of a condensation relationship might cause two vertices to now have the same support. If such a situation is detected, the edge between these two vertices is removed from the component.
(b) Put in a set the p-invariants from the source (target) vertices that has only one p-invariant. These p-invariants are the core source (target) p-invariants. If present, remove the target p-invariants of the previous exploration step subgraph from this exploration step’s target p-invariants.
(c) Remove the core source p-invariants from the target p-invariants and the core target p-invariants from the source p-invariants. If there is no p-invariant left in the remaining target p-invariants, this might indicate a modeling assumption that is not modeled explicitly. Send a request to the user to identify the correct interaction partner.
(d) In the sets of source and target p-invariants, for each partition keep only the sets with the lowest cardinality to keep the minimal covering sets of p-invariants for both partitions. If there are still more than one p-invariant each in the source and target sets, remove from either set the intersection between source and target sets.
(e) Influence arcs are then created (but not added to the influence graph yet) between all remaining sources and target p-invariants. The quadruplets from the initial interaction arcs are associated with the new influence arcs.

The **fourth processing step** of this rule identifies which influence arcs among those created at the precedent step are a signal condensation relationship and if a recursive association with previous relationships is necessary. Because they only have one target, arcs representing enzymatic reactions do not condense the signal. For the remaining arcs, a signal condensation relationship is set between the source p-invariant and its target: the source p-invariant becomes a codomain of the target p-invariant and the target p-invariant becomes a domain of the source p-invariant. For molecular complexes with more than two partners, condensation relationship must be established recursively. To decide if this recursion is necessary, it is verified if the source place of the arcs’ quadruplets contains p-invariants that are already codomains of the source of the arc. When this condition is met, two recursive updates are performed: first, the other codomains of the source p-invariant become codomains of the target p-invariant and it becomes their domain; second, other domains of the target p-invariant become domains of the source p-invariant and it becomes their codomain. All these condensation relationships are used in the second processing step to condense signals in vertices in the next iterations of this procedure.

Finally, **the fifth processing step** is to add the new influence arcs from this exploration step to the influence graph, condensing and non-condensing arcs alike. The arcs’ quadruplets are validated to remove pairs of quadruplets involving transitions forming trivial t-invariants. If the vertices of an arc are absent from the graph, they are created. If the two vertices of an arc exist and an influence arc is already between them in the graph, the quadruplets of the new arc are added to the quadruplets of the existing arc. Finally, the p-invariant targets of the new arcs are stored for the operation 3b of the next iteration. This completes the processing of the current exploration step subgraph. The processing of the subgraph for the next step will begin with probing each vertex for a signal condensation relationship: if in the same vertex, one p-invariant is the codomain of another, then the codomain p-invariant is removed from the vertex. These processing steps are performed on every subgraph. Establishing condensation relationships is the most important part of this rule. At each step, these relationships are used to remove some p-invariants of the vertices of the interaction graph, leaving only the p-invariants that have direct interactions, which then become influence arcs.

*Example 11:* Figure 8 presents five simple cases of interaction edges between source and target vertices converted into influence arcs with the five processing steps. In Figure 8a, there are two core p-invariants and the transformation results simply into the influence arc *y*_1_ → *y*_2_. The following condensation relationship is established if this edge is not associated with an enzymatic reaction: *y*_1_ becomes a codomain of *y*_2_ and *y*_2_ becomes a domain of *y*_1_. In Figure 8b, *y*_1_ is a core p-invariant of the source vertex because it is a single p-invariant. It is thus removed from the target vertex. This also results in the influence arc *y*_1_ → *y*_2_. In Figure 8c, *y*_2_ is a core p-invariant of the target vertex and is removed from source vertex. This results in the influence arc *y*_1_ → *y*_2_. In Figure 8d, the two interaction edges are evaluated one after the other. In the first evaluation, the top two vertices are identical to Figure 8b and generate the influence arc *y*_1_ → *y*_2_. In the evaluation of the second edge, the first step is to verify if the signal should be condensed using existing condensation relationships. Because *y*_1_ has been identified as a codomain of *y*_2_ at a previous step, the p-invariant *y*_1_ is removed from the vertices. This results in the influence arc *y*_2_ → *y*_3_. *y*_2_ becomes a codomain of *y*_3_ and *y*_3_ becomes a domain of *y*_2_. Through the recursive update, *y*_1_ becomes a codomain of *y*_3_. In Figure 8e, *y*_1_ is a core p-invariant of the source vertex and is removed from the target vertex and two p-invariants remain. A request is sent to the user to validate the target: *y*_2_, *y*_3_ or both. One or two influence arcs will be created accordingly.

**Figure 8:**
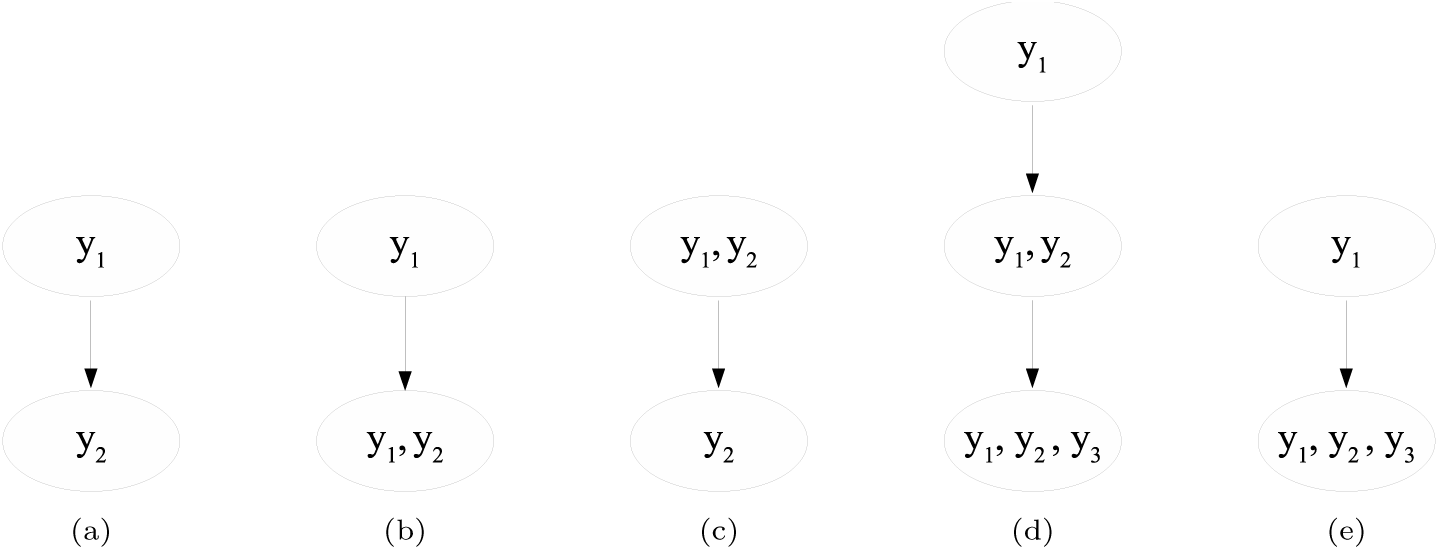
Source and target vertices with edges from an interaction graph to be converted into an influence graph. (a), (b) and (c) are converted into the influence arc *y*_1_ *→ y*_2_ because in each case, the core source vertex is *y*_1_ and the core target vertex is *y*_2_, (d) is converted into the influence arcs *y*_1_ *→ y*_2_ *→ y*_3_ because *y*_3_ is the only core target remaining for the second edge after the elimination of *y*_1_ and *y*_2_ due to a signal condensation relationship established by the first edge. In (e), because more than one p-invariant remains in the target vertex after analysis, the user is prompted to choose if an influence arc is established from *y*_1_ to *y*_2_, from *y*_1_ to *y*_3_ or both.

*Example 12:* To exemplify the rule for building the influence graph, we introduce a Petri net model more complex than *N*_1_. For this example, we use the Petri net *N*_2_ shown in Figure 9 and perform the first two phases of the algorithm to transform *N*_2_ into its influence graph representation. This model represents the molecular binding of three molecules to form a complex. Molecule *A* is a ligand and can bind to molecule *B*, a membrane receptor, to form the complex *AB*. Molecule *B* can also bind to molecule *C*, a small switch-like protein, to form the complex *BC*. These bindings are not mutually exclusive and can happen in any order to form the trimeric complex *ABC*, a ligand-receptor-protein complex. Every binding reaction has its unbinding reverse counterpart. The formation of the trimer *ABC* leads to the activation of the switch-like protein in the form of *C_prime_*. At the beginning of the preparatory phase, the p-invariant analysis on this model detects three p-invariants, *y_A_*, *y_B_* and *y_C_* for the three molecular types, and produces the lookup table *placeToInvariant* (Table 7). No pairs of p-invariants from the model are considered similar. The three p-invariant subnets *N*_2_*_A_*, *N*_2_*_B_* and *N*_2_*_C_* are analyzed for signaling segments. Three signaling segments are identified: 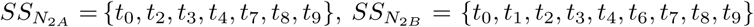, 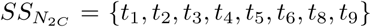. The Petri net model is explored from place *p*_0_ (molecule *A*), identified as the source of the biological signal by the user. The exploration of the Petri net produces the lookup table *RFAP* with the quadruplets that might become interaction edges (Table 8). The first rule of the second phase of the algorithm creates the interaction graph shown in Figure 10: six vertices are created for the different sets *E_p_* from the lookup table *placeToInvariant* and eighteen edges connect the vertices, listed with their associated quadruplet in Table 10. No edge was created for the quadruplet [*p*_6_*, p*_4_*, t*_5_, 2] since its source and target places are associated with the same *E_p_*: *y_C_*. This rule also generates the lookup table *OE* (Table 9). The next rule uses signaling segments to eliminate some edges from the interaction graph. Eleven edges are removed (listed in red in the Table 10), notably every edge from the exploration step 2. The resulting graph is shown in Figure 11.

**Figure 9:**
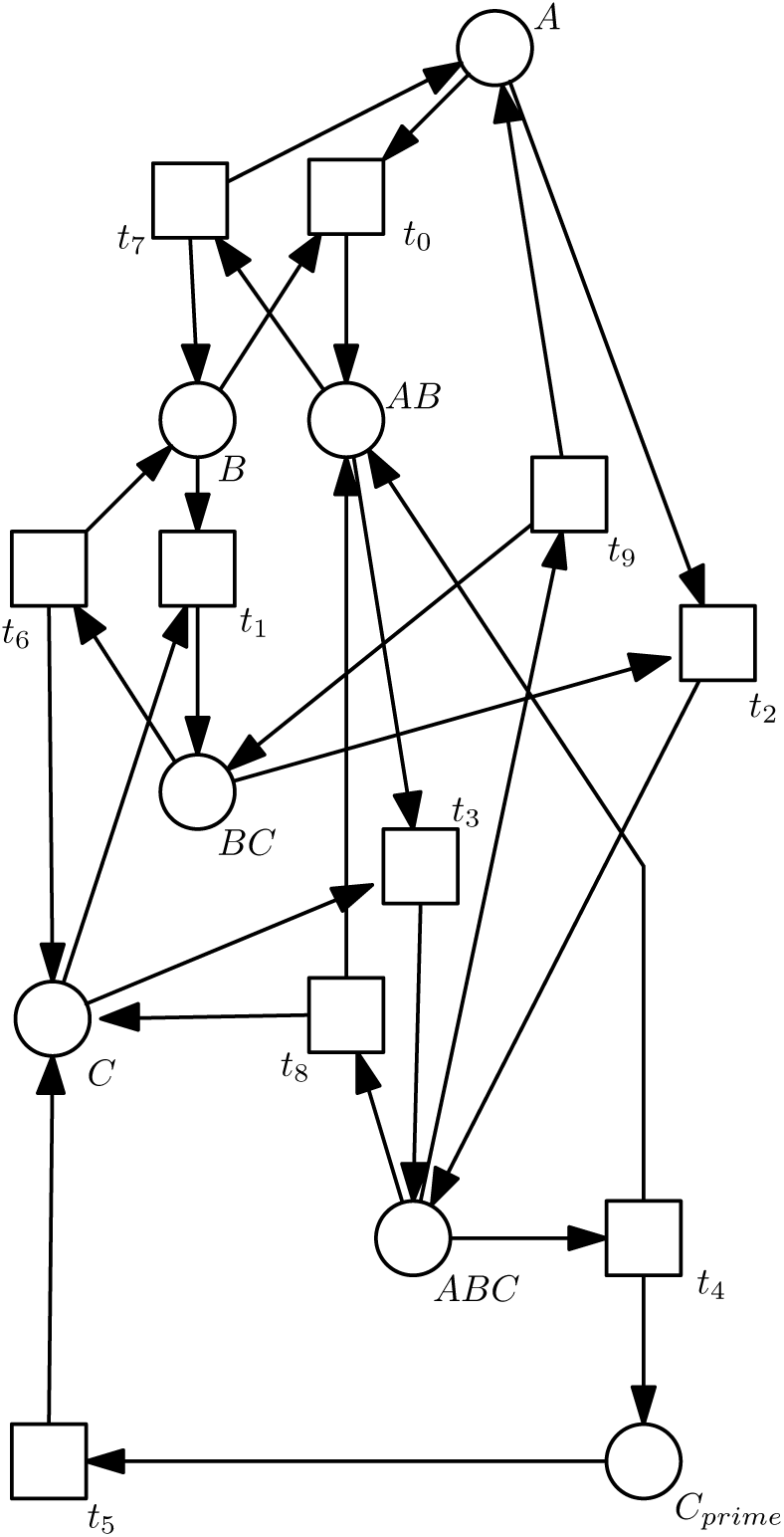
The Petri net *N*_2_ represents several binding reactions creating the trimer with the molecules A, B and C. Once in the trimer ABC, the molecule C is activated in the form of Cprime through transition *t*_4_. In this signaling pathway, the source of the signal is molecule A (place A).

**Table 7:**
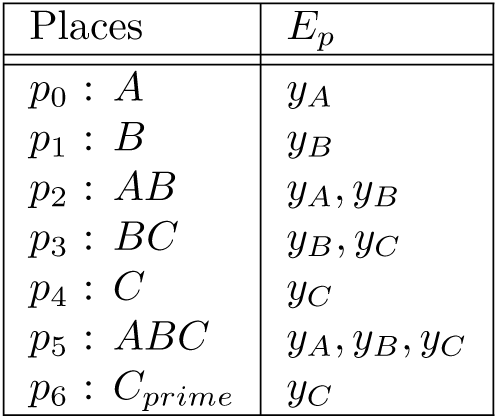
The lookup table *placeT oInvariant* after the p-invariants analysis, which associates each place of the Petri net *N*_2_ to the set of p-invariants the place belongs to.

**Table 8:** The lookup table *RF AP* after the exploration of the Petri net *N*_2_ from the source place.

| Places | Quadruplets |
| --- | --- |
| $p_0$ | $[p_0, p_2, t_0, 0], [p_0, p_5, t_2, 0]$ |
| $p_1$ | $[p_1, p_3, t_1, 2], [p_1, p_2, t_0, 2]$ |
| $p_2$ | $[p_2, p_5, t_3, 1], [p_2, p_1, t_7, 1], [p_2, p_0, t_7, 1]$ |
| $p_3$ | $[p_3, p_5, t_2, 2], [p_3, p_1, t_6, 2], [p_3, p_4, t_6, 2]$ |
| $p_4$ | $[p_4, p_5, t_3, 2], [p_4, p_3, t_1, 2]$ |
| $p_5$ | $[p_5, p_4, t_8, 1], [p_5, p_3, t_9, 1], [p_5, p_2, t_4, 1],$<br>$[p_5, p_2, t_8, 1], [p_5, p_6, t_4, 1], [p_5, p_0, t_9, 1]$ |
| $p_6$ | $[p_6, p_4, t_5, 2]$ |

**Figure 10:**
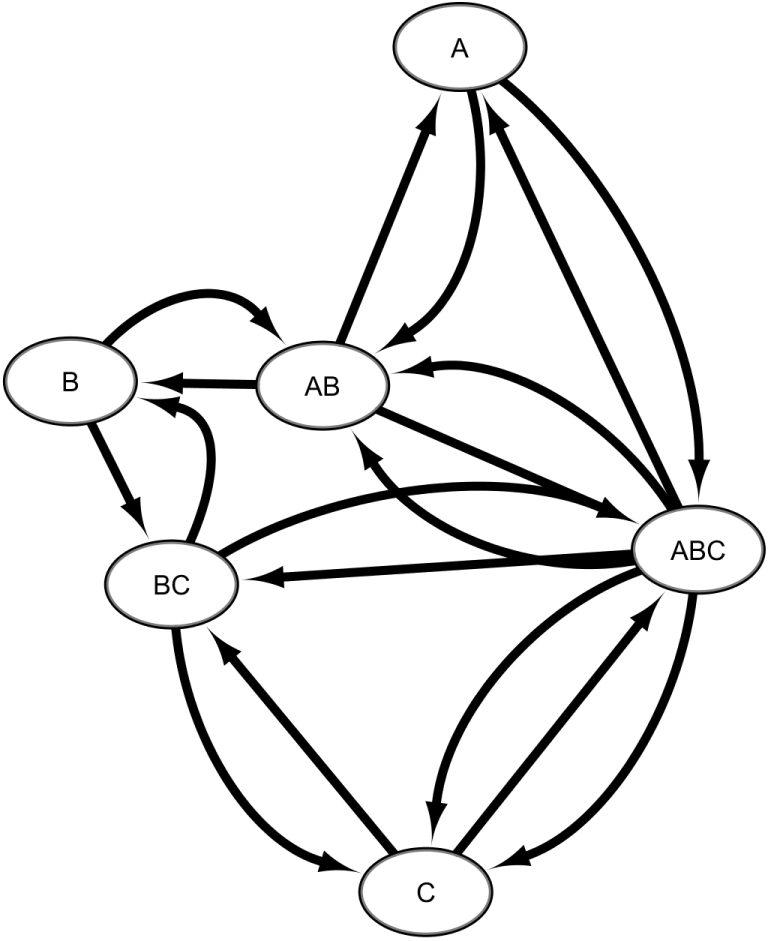
Interaction graph built from the Petri Net *N*_2_ after the execution of the first rule of the second phase with its vertices and their associated p-invariants, and its edges, which associated quadruplets are listed in Table 10.

**Table 9:** The lookup table *OE* for the Petri net *N*_2_, representing each step and their associated quadruplets. Each quadruplet corresponds to an egde in the interaction graph shown in Figure 10.

| Step | Quadruplets |
| --- | --- |
| 0 | $[p_0, p_2, t_0, 0], [p_0, p_5, t_2, 0]$ |
| 1 | $[p_5, p_4, t_8, 1], [p_5, p_3, t_9, 1], [p_5, p_2, t_4, 1],$<br>$[p_2, p_5, t_3, 1], [p_5, p_2, t_8, 1], [p_2, p_1, t_7, 1],$<br>$[p_5, p_6, t_4, 1], [p_5, p_0, t_9, 1], [p_2, p_0, t_7, 1]$ |
| 2 | $[p_4, p_5, t_3, 2], [p_1, p_3, t_1, 2], [p_1, p_2, t_0, 2],$<br>$[p_3, p_5, t_2, 2], [p_4, p_3, t_1, 2], [p_3, p_1, t_6, 2],$<br>$[p_3, p_4, t_6, 2]$ |

**Table 10:** List of edges in the interaction graph for the Petri Net *N*_2_, before the signaling segments rule is applied and after. Every edge marked in red is removed by the rule.

| Source vertex | Target vertex | Quadruplets |
| --- | --- | --- |
| $\{A\}$ | $\{A, B\}$ | $[p_0, p_2, t_0, 0]$ |
| $\{A\}$ | $\{A, B, C\}$ | $[p_0, p_5, t_2, 0]$ |
| $\{B\}$ | $\{A, B\}$ | $[p_1, p_2, t_0, 2]$ |
| $\{B\}$ | $\{B, C\}$ | $[p_1, p_3, t_1, 2]$ |
| $\{C\}$ | $\{A, B, C\}$ | $[p_4, p_5, t_3, 2]$ |
| $\{C\}$ | $\{A, B\}$ | $[p_4, p_3, t_1, 2]$ |
| $\{A, B\}$ | $\{A\}$ | $[p_2, p_0, t_7, 1]$ |
| $\{A, B\}$ | $\{B\}$ | $[p_2, p_1, t_7, 1]$ |
| $\{A, B\}$ | $\{A, B, C\}$ | $[p_2, p_5, t_3, 1]$ |
| $\{B, C\}$ | $\{B\}$ | $[p_3, p_1, t_6, 2]$ |
| $\{B, C\}$ | $\{C\}$ | $[p_3, p_4, t_6, 2]$ |
| $\{B, C\}$ | $\{A, B, C\}$ | $[p_3, p_5, t_2, 2]$ |
| $\{A, B, C\}$ | $\{A\}$ | $[p_5, p_0, t_9, 1]$ |
| $\{A, B, C\}$ | $\{A, B\}$ | $[p_5, p_2, t_8, 1]$ |
| $\{A, B, C\}$ | $\{A, B\}$ | $[p_5, p_4, t_4, 1]$ |
| $\{A, B, C\}$ | $\{B, C\}$ | $[p_5, p_3, t_9, 1]$ |
| $\{A, B, C\}$ | $\{C\}$ | $[p_5, p_6, t_4, 1]$ |
| $\{A, B, C\}$ | $\{C\}$ | $[p_5, p_4, t_8, 1]$ |

**Figure 11:**
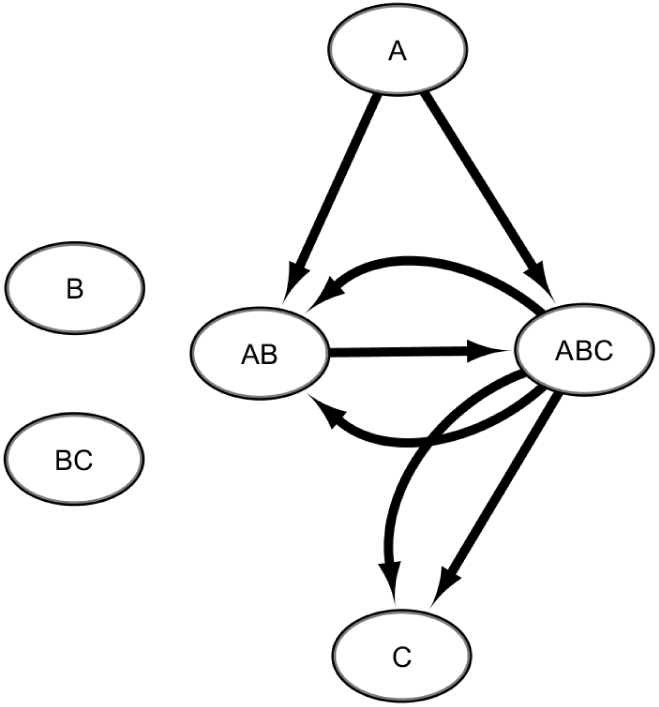
Interaction graph for the Petri Net *N*_2_ with the remaining edges after applying the signaling segments rule when the place *A* is considered the source of the signal.

To conclude the second phase of the algorithm, the influence graph is built one exploration step at a time. For step 0, a subgraph is extracted. It contains the three vertices and the two edges associated with step 0 in *OE*: A → AB and A → ABC. (processing step 1). At this first iteration, no signal condensation relationship exists yet, so there is no signal to condense (processing step 2). This subgraph is a maximally bipartite component by itself (processing step 3). The supports of the three vertices are all different, so no edge has to be removed (step 3a). The p-invariant A is identified as a core source p-invariant, and there is no core target p-invariant since there is no set with only one p-invariant in the target vertices AB and ABC (step 3b). The core p-invariant is removed from the other partition vertices: A is removed from the vertices AB and ABC, which become B and BC (step 3c). A is identified as the source, and B, being the set with the lowest cardinality, is identified as the target; since source and target now have only one p-invariant, no further processing is needed (step 3d). This result shows that A interacts directly with B but indirectly with C since C is never in complex only with A (thus without B). An influence arc between vertices A and B is created and the quadruplets [*p*_0_*, p*_5_*, t*_2_, 0] and [*p*_0_*, p*_2_*, t*_0_, 0] from the two interaction edges are added to this new arc. This arc is identified as a condensing one, and a condensation relationship is established: A is a codomain of B and B is a domain of A (processing step 4). The two new vertices A and B and their connecting arc are then added to the influence graph (processing step 5).

For exploration step 1, another subgraph is extracted: it contains the vertices AB, ABC and C and their five connecting arcs (processing step 1). With the previously established signal condensation relationship (A is a codomain of B), the signal is condensed in the vertices of this subgraph: the p-invariants A is removed from the two vertices in which the p-invariant B is also present (processing step 2). This subgraph has two maximal bipartite components: the first component is B → BC and the second component has three vertices and the four edges BC → B (twice), BC → C (twice) (processing step 3). In the first component, all the vertices’ supports are different, so no edge has to be removed (step 3a). The core source and target p-invariants are B and C respectively (step 3b) and no further processing occurs for the steps 3c and 3d. An influence arc is created between vertices B and C with the quadruplet [*p*_2_*, p*_5_*, t*_3_, 1] (step 3e). In the second maximal bipartite component, all the supports of the vertices are also different, so no edge has to be removed (step 3a). There is no core source p-invariant in BC, but C is a core target p-invariant as B is removed since it is the target of the iteration of the previous step (step 3b). The core target p-invariant C is removed from the source p-invariants, leaving B (step 3c). As a result, B is the sole source p-invariant and C is the sole target p-invariant (step 3d). An influence arc is created between vertices B and C with the two quadruplets [*p*_5_*, p*_6_*, t*_4_, 1] and [*p*_5_*, p*_4_*, t*_8_, 1] associated with the edges linking these vertices in the subgraph (step 3e). At processing step 4, the two new influence arcs for the exploration step 1 are not enzymatic reactions and thus condense the signal, and a condensation relationship is established: B is a codomain of C and C is a domain of B. Since B already has a codomain, another condensation relationship between A and C is recursively established: A is a codomain of C and C is a domain of A. At the processing step 5, the two new arcs are candidates to the influence graph: the addition of the first arc causes the creation of the vertex C in the influence graph; for the second arc, an arc between the vertices B and C now exists and consequently, its quadruplets are tentatively added to the existing arc. Since the transitions *t*_3_ and *t*_8_ are the support of a trivial t-invariant in the Petri net, the two quadruplets containing them are discarded and only the quadruplet [*p*_5_*, p*_6_*, t*_4_, 1] remains. There is no more subgraph to analyze since every edge from exploration step 2 has been previously removed by the signaling segments rule. The resulting influence graph for the Petri net *N*_2_ is shown in Figure 12. Table 11 details the influence arcs and their associated quadruplets. It is interesting to note that the transitions of the quadruplets of the arc between vertices A and B are *t*_0_ and *t*_2_, which correspond to the activation of B by the binding of A, and that the transition of the quadruplet of the arc between vertices B and C is *t*_4_, which correspond to the activation of C following the formation of the trimer ABC. These transitions are key to the biochemical process modeled by the Petri net *N*_2_.

**Figure 12:**
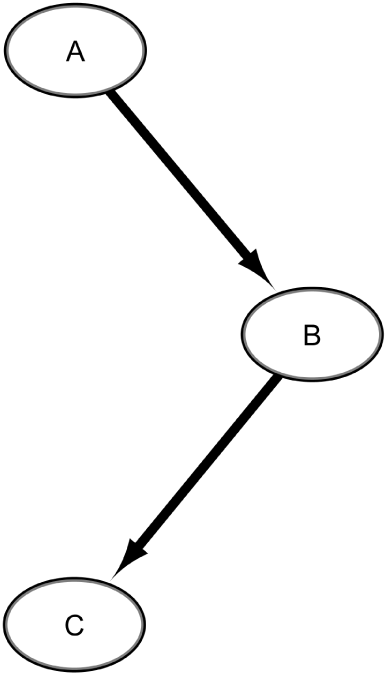
Influence graph created for the Petri Net *N*_2_ after processing the interaction graph shown in Figure 11 with the third rule of the second phase. The quadruplets of the graph arcs are listed in Table 11.

**Table 11:** The arcs of the influence graph (Figure 12) and their associated quadruplets, for the Petri Net *N*_2_.

| Source Vertex | Target Vertex | Quadruplets |
| --- | --- | --- |
| A | B | $[p_0, p_5, t_2, 0], [p_0, p_2, t_0, 0]$ |
| B | C | $[p_5, p_6, t_4, 1]$ |

The second phase of the transformation algorithm ends with the creation of the influence graph, which will be annotated in the next phase. The three transformation rules of this phase are described summarily in Algorithm 2, with the removal of interaction edges by the signaling segments rule detailed in Algorithm 2.1 and the building of the influence graph detailed in Algorithm 2.2 where the instructions for the processing steps 1 to 5 are clearly indicated. The algorithm of the second phase is provided with more details in Algorithm S2.

##### Algorithm 2 Building the influence graph from the interaction graph (phase II)

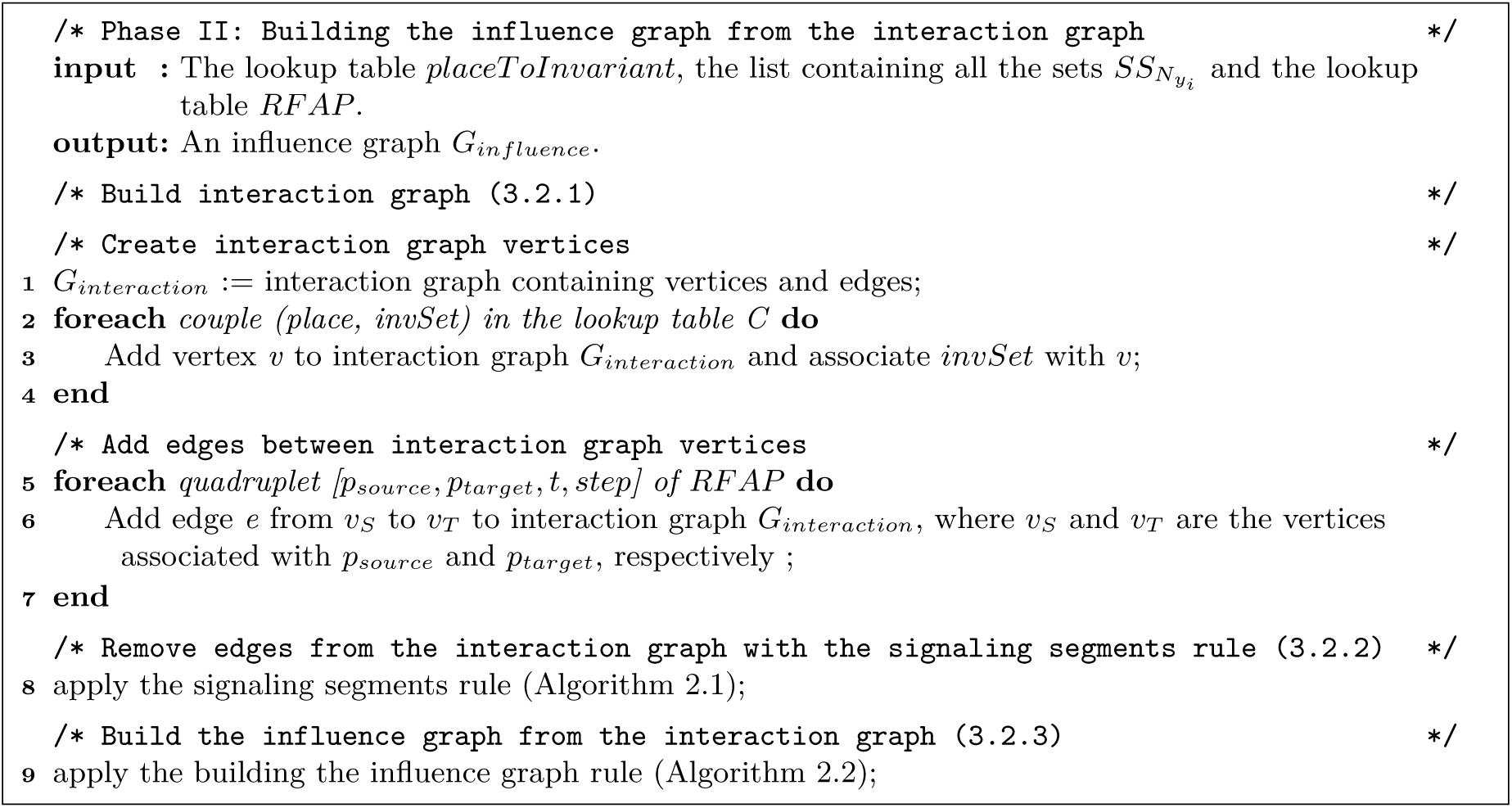

##### Algorithm 2.1 Algorithm for removing interaction edges with the signaling segments rule

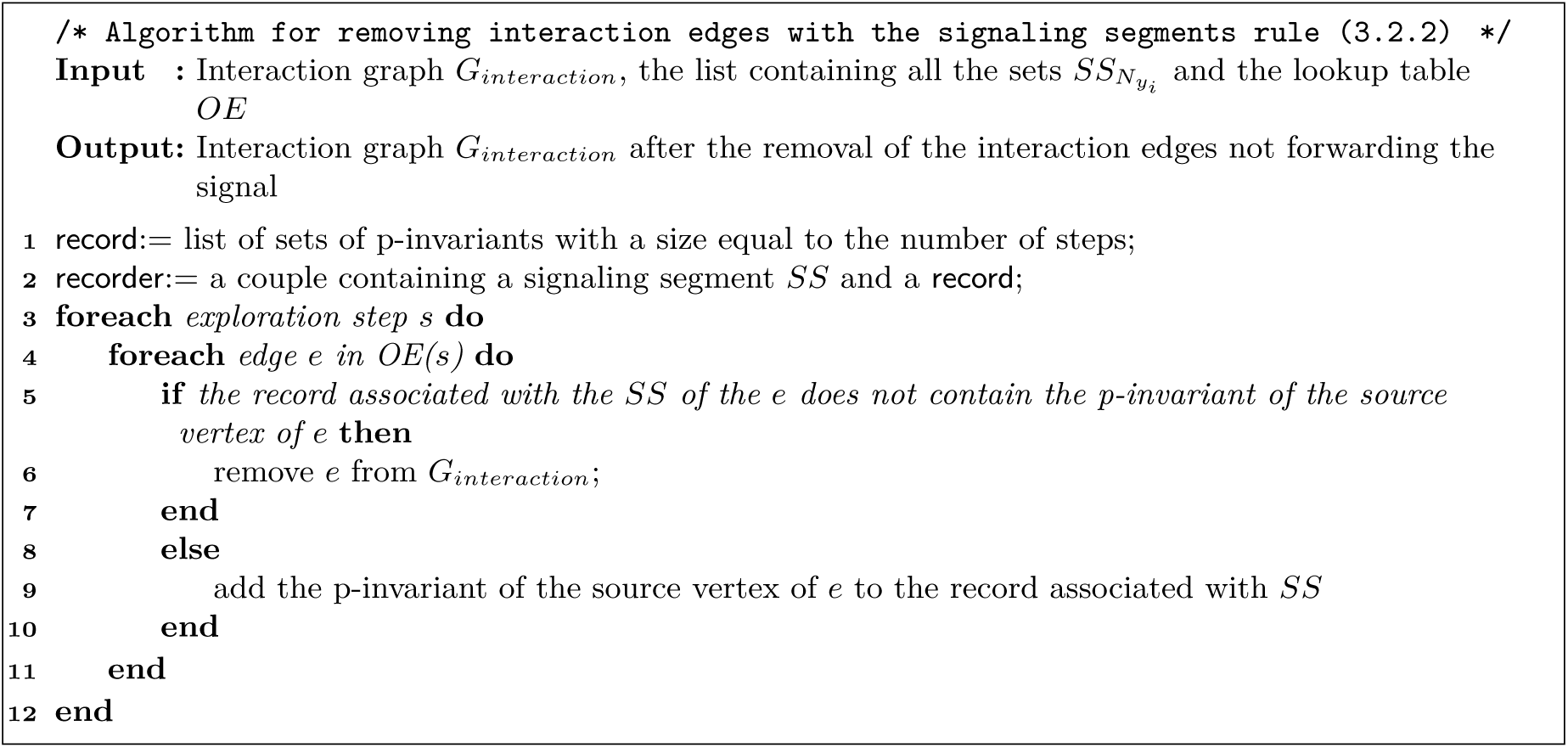

### 3.3. Phase III: Annotating the influence graph and determining the type of influence of the arcs

In the second phase of the transformation algorithm, the structural properties of a Petri net model and complementary modeling information were used to build the model’s representation as an influence graph. In the final phase, further structural information is used to determine the type of influence for the arcs of the graph. Three transformation rules are necessary to do so. First, places that participate in the transmission of the signal downstream in the signaling pathway are identified as active molecular forms in the model. These places are labeled signaling places. Similarly, transitions that transmit the signal by activating or inactivating a downstream molecule are identified by the second rule. Finally, this information allows for the assignment of an influence type (activation or inhibition) to the arcs of the influence graph.

#### Algorithm 2.2 Algorithm for building the influence graph.

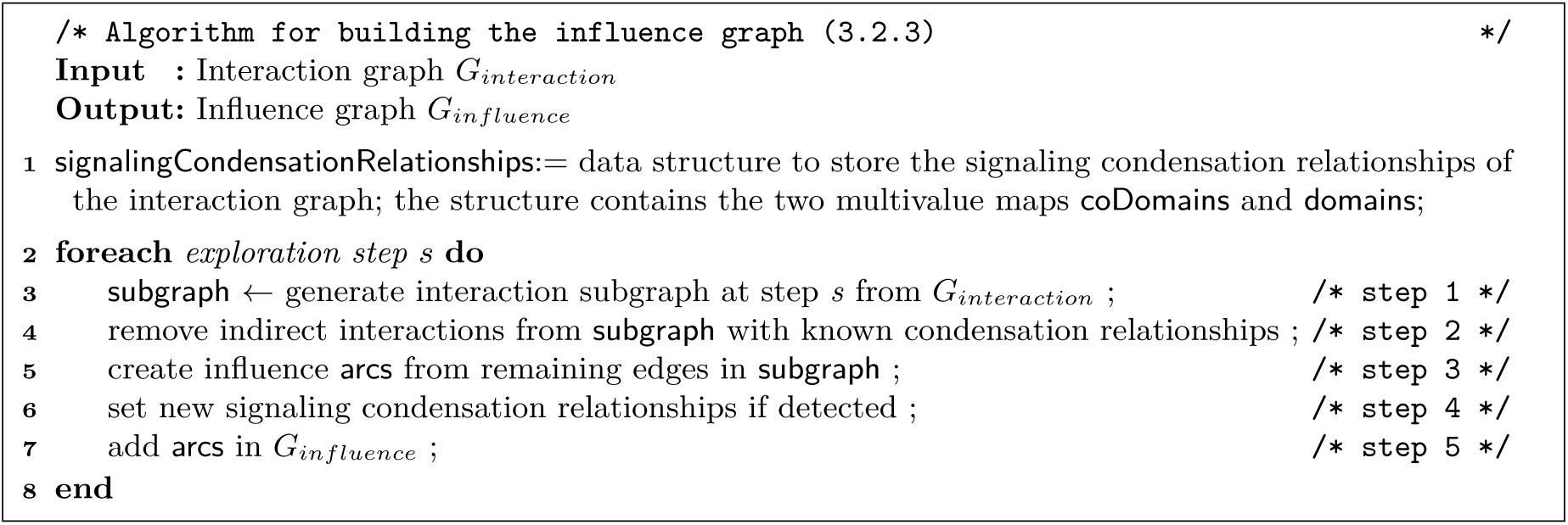

#### 3.3.1. Detecting signaling places and associating them with the vertices of the influence graph

After creating vertices and connecting them with arcs, the influence graph of a signaling model can now be annotated with additional dynamic information from the model. The different configurations of a molecular species, which correspond to the different places of the Petri net, can be associated with either the active or the inactive state of the species. When linked to the active state, the places are detected by this rule as signaling places, otherwise they are non-signaling places. Consequently, the support of every p-invariant of the Petri net model is divided into two subsets: the signaling and non-signaling places.

To identify the places that correspond to the active state, we propose two conditions in the following criteria: the signaling places of a vertex (by default, these places are in the support of this vertex) are either a source place in the quadruplet of an arc connecting this vertex to a target vertex or a target place in a similar quadruplet. In this latter condition, the places must also be in the support of the target vertex. This criterion is formally defined in definition 19. For example, the place corresponding to an enzyme that catalyzes a reaction is detected as a signaling place according to the first condition of the criteria. Another example is the activation of species B by species A through the binding of two molecules. The place associated with molecule A is detected as a signaling place according to the first condition, and the place associated with molecular complex AB is detected as a signaling place according to the second condition. In signaling models, these two signal transduction mechanisms are the most common. A less frequent mechanism – an active biochemical signal causing the unbinding of a complex – is not covered by this definition and requires a specific operation.

*Definition 19:* Let *v_a_* and *v_b_* be two vertices in an influence graph and *e*(*v_a_*, *v_b_*) an arc from *v_a_* to *v_b_*. The sets ∥*v_a_*∥ and ∥*v_b_*∥ are the supports of *v_a_* and *v_b_*, respectively. Let *p* be a Petri net place where *p* ∈ ∥*v_a_*∥. The place *p* is considered as signaling for the vertex *v_a_* if ∃ quadruplet *q*_1_ [*p, p_target_, t, step*] such as *q*_1_ ∈ *Tr*-*G_influence_*(*e*) or if ∃ quadruplet *q*_2_ [*p_source_, p, t, step*] such as *q*_2_ ∈ *Tr*-*G_influence_*(*e*) and *p* ∈ ∥*v_b_*∥.

This definition to detect the active forms of a signaling molecular species relies on the connectivity of a vertex to its targets in the influence graph. As a result, it cannot be applied to the vertices without any target, such as the vertices that are leaves in a graph. The first operation of this rule is thus to find the leaves of the influence graph. Then a verification of the edges connecting the leaves to their source vertices is performed. Leaves are signaling endpoints of the modeled pathways, however leaves in the influence graph so far might be covalent inhibitors in some models. Covalent inhibitors act as buffers for other activated signaling molecules. In this case, the correct biochemical influence is from the inhibitor to the inhibited molecule, and the user is prompted to provide this information, which is used to correct this edge and its quadruplets if necessary.

The second operation of this rule is to identify the signaling places of each vertex with the general criteria of Definition 19. Thirdly, five specific situations are detected and processed. For leaf vertices, the user is prompted to identify the signaling places among the vertex support. When the signal causes the unbinding of a complex (a situation detected by an edge being associated with a quadruplet in which a transition has one preplace and two or more postplaces), the user is prompted to identify the signaling places of the source vertex of this edge. If for a vertex every place is signaling or every place is non-signaling, the user is prompted to choose the signaling places manually. This situation occurs when the inactive form of a molecular species can signal to another downstream species at a basal level of activity. Finally, all the places of the vertex that is the signaling source of the model are labeled as signaling places.

When the vertices of the influence graph have been processed by this rule, each vertex is associated with a set of signaling places, which is a subset of its support. The remaining places are considered non-signaling.

#### 3.3.2. Associating signaling transitions to influence arcs

During the construction of the influence graph in rule 3.2.3, one or multiple quadruplets are associated to each influence arc of the graph. Not all of these quadruplets contain a transition that conveys a biological signal of activation or inhibition. This rule will identify the quadruplets containing a transition that corresponds to the activation or inactivation of a downstream molecule and remove the others.

*Definition 20:* Let *v_a_* and *v_b_* be two vertices of *G_influence_* and *e*(*v_a_*, *v_b_*) the influence arc from *v_a_* to *v_b_*. Let [*p_source_, p_target_, t, step*] be a quadruplet *q* such as *q* ∈ *Tr*-*G_influence_*(*e*). If *p_source_*∈*/* signaling places of *v_a_* and neither *p_target_*∈ signaling places of *v_b_* nor there is no preplace of *t* (^•^*t*) ∈ signaling places of *v_b_*, then *q* is removed from *Tr*-*G_influence_*(*e*).

This definition detects the following two conditions: if the source place of a quadruplet is a signaling place of the source vertex, or if the target place of the same quadruplet or a preplace of the transition is a signaling place of the target vertex. In other words, this corresponds to molecular reactions in which an active molecular species activates or deactivates another one. The algorithm iterates over every quadruplet of the arcs and applies these conditions. After the execution of this rule, the arcs of the influence graph are associated only with quadruplets that can represent either a biological activation or an inhibition. The other quadruplet or discarded. The next rule will identify the type of influence and label the arcs accordingly.

#### 3.3.3. Determining the type of the arcs of the influence graph

In an influence graph, the arcs can have a type. So far in the transformation algorithm, no influence type has been assigned to the arcs of the graph built from the biological signaling model. As a biochemical signal is transduced in a cell through molecular reactions, molecules are activated or deactivated through binding/unbinding and enzymatic reactions. In a previous rule (Section 3.3.1), places were identified as signaling or non-signaling places, which can also be construed as on and off states. Consequently, an activation influence should turn on a molecular species by switching some places from non-signaling to signaling and an inhibitory influence should do the opposite.

*Definition 21:* Let *v_a_* and *v_b_* be two vertices of *G_influence_* and *e*(*v_a_*,*v_b_*) the influence arc from *v_a_* to *v_b_*. The type of arc *e* is inhibitory if ∃ a quadruplet *q* [*p_source_*, *p_target_*, *t*, step] such as *q* ∈ *Tr*-*G_influence_*(*e*) in which *p_target_* is a non-signaling place of ||*v_b_*|| (the support set of the p-invariant associated to *v_b_*) and the set of pre-places •*t* except *p_source_* contains only signaling places of ||*v_b_*||.

In the algorithm, every arc is initially labeled as an activation arc in the influence graph. Then, the algorithm iterates over each arc edge and tests the condition in Definition 21, which verifies if the state of a molecular species is switched from signaling places to non-signaling through the transition of at least one quadruplet in *Tr*-*G_influence_*(*e*). If the condition is fulfilled, then the type of the arc is changed to inhibitory. At the end of this rule, the transformation from a Petri net biological model to its annotated representation as an influence graph is complete.

The algorithm for the three rules of this phase for the annotation of the influence graph is detailed in Algorithm 3. The algorithm is also provided in a version with more formal details in Algorithm S3. In the next section, we present an example of the creation of an influence graph from a Petri net biological model through the three phases of the transformation algorithm.

##### Algorithm 3 Annotating the influence graph

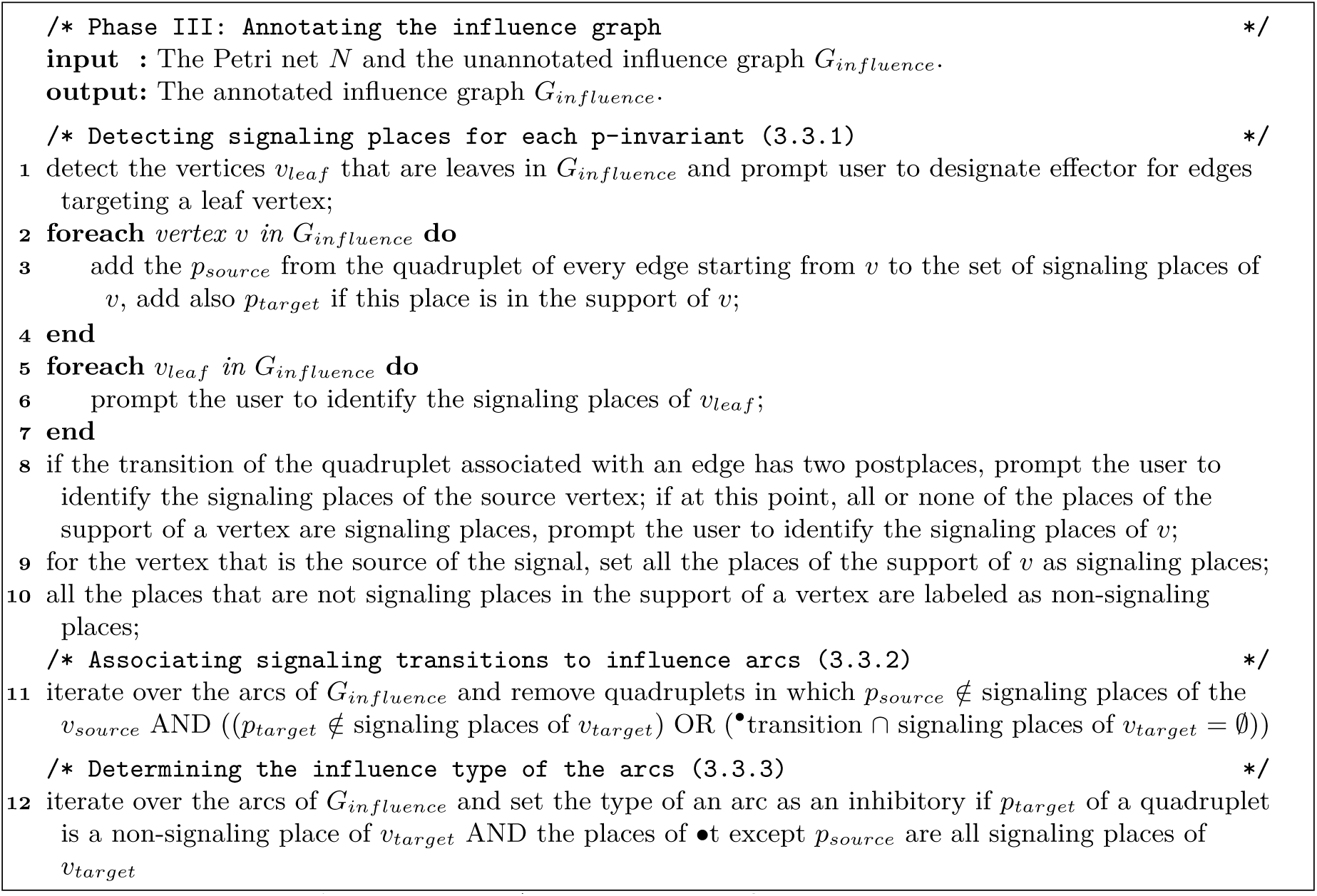

## 4. Example of the execution of the algorithm: transformation of the Petri net model of the GPCR-PKA-MAPK signaling network into an annotated influence graph

In this section, we transform the Petri net representation of the differential equations model of the signaling pathway from the beta-adrenergic receptor (BAR) to the kinase MAPK1,2 through the cAMP/PKA/B-Raf/MAPK1,2 network in neurons [38] into an annotated influence graph using the transformation algorithm presented in this paper. This model contains only mass-action reactions and one-step enzymatic reactions. This transformation will highlight the signaling flow in the model and its regulatory mechanisms.

The agonist that activates the G-protein-coupled receptor of this network is isoproterenol, a molecular analog of epinephrine. This receptor can bind to a G protein to form a molecular complex. In the complex, the presence of the ligand activates the G protein, a heterotrimer in itself, and the alpha subunit (G*_α_*_s_) that is now bound to GTP instead of GDP is separated from the beta/gamma subunits (G*_βγ_*). The hydrolysis of GTP to GDP enables the return of the inactivated alpha subunit back into the complex with the beta/gamma subunit. When the separated active G*_α_*_s_ binds to the enzyme adenylyl cyclase, it increases the enzyme conversion of adenosine triphosphate (ATP) into the second messenger cyclic adenosine monophosphate (cAMP). Once separated from the alpha subunit, G*_βγ_* can bind to a G-protein-coupled receptor kinase (GRK), an enzyme that can phosphorylate the receptor, thus regulating the receptor sensitivity through inhibition.

The two regulatory subunits of protein kinase A (PKA) together have four binding sites for cAMP. Once the four sites are occupied by the second messenger, the catalytic subunits of PKA are activated and free to phosphorylate the targets of this enzyme. A first target of PKA is the kinase B-Raf, which activates the MEK-MAPK cascade. A second target is the protein tyrosine phosphatase (PTP), which deactivates MAPK through dephosphorylation. The last target of PKA is the phosphodiesterase PDE4 which catalyzes the hydrolysis of cAMP to AMP. As a result of all these interactions, there are three regulatory network motifs in this signaling pathway: a negative feedback loop between BAR, G*_βγ_* and GRK; a second negative feedback loop formed by the reactions between cAMP, PKA and PDE4; and a coherent feedforward loop with PKA as its source, MAPK as its target and B-Raf/MEK/MAPK and PTP being the two paths linking the two molecular species.

Figure 13 shows the Petri net *N_GP CR_* of this reaction-based model. It has 33 places and 43 transitions. There are only two types of reaction in this model: binding reactions and enzymatic reactions. Read arcs connect enzyme places and their reaction transitions. We now apply the three phases of the transformation algorithm and present the intermediate data structures and graphs, and in the end the annotated influence graph.

**Figure 13:**
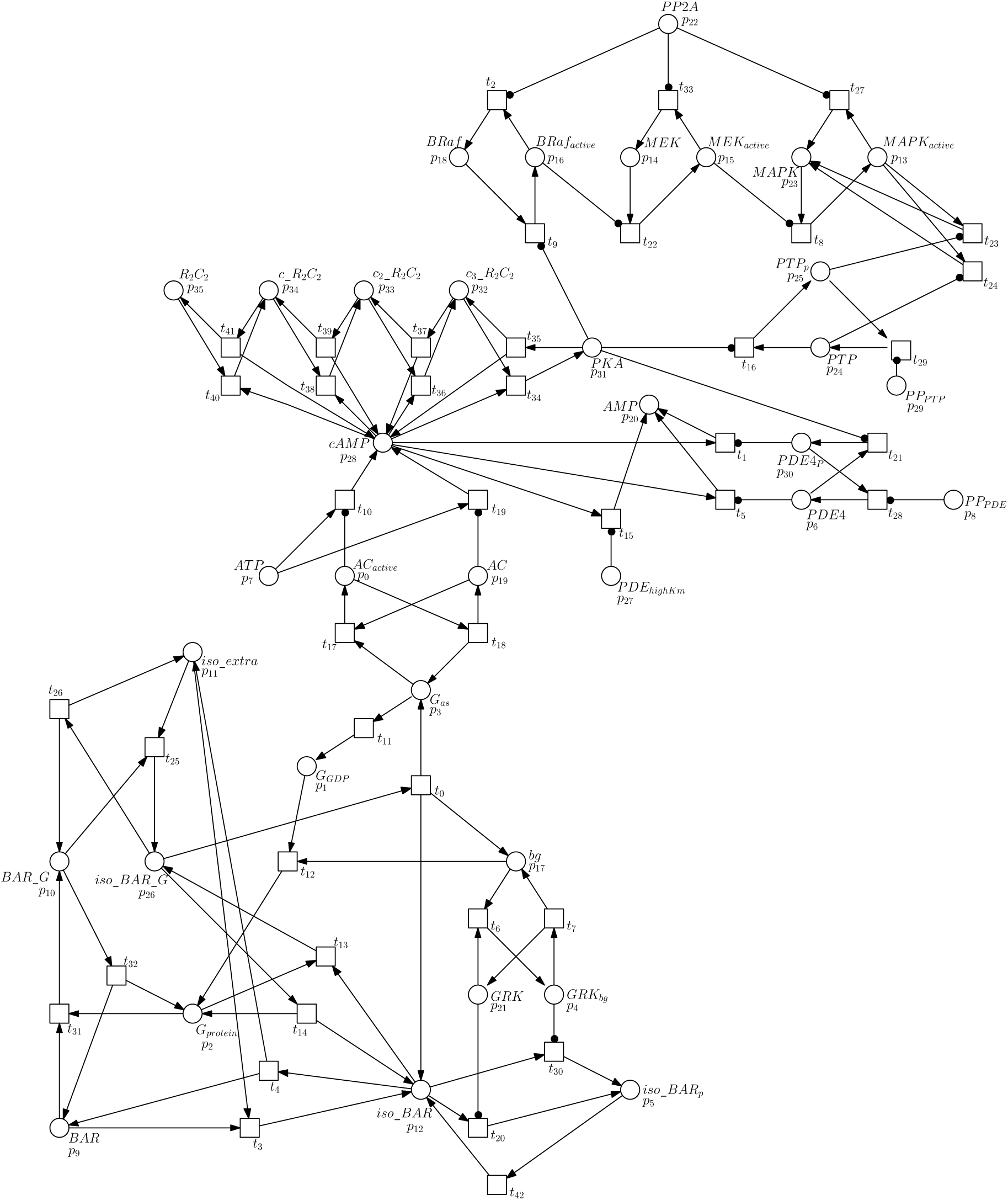
Petri net model *N_GPCR_* of the signaling network from the beta-adrenergic receptor to PKA/MAPK

### 4.1. Phase I: Generation of the required data structures from an analysis of the Petri net model of the GPCR-PKA-MAPK signaling network

The three preparatory operations of the first phase of the algorithm are applied to the Petri net *N_GP CR_*, resulting in the lookup table *placeToInvariant*, the list of the signaling segments of the model and the lookup table *RFAP*.

#### 1. P-invariant analysis and validation

A p-invariant analysis is performed on the Petri net *N_GP CR_*, and 17 p-invariants are found. The lookup table *placeT oInvariant* is created (Table S1) where each place of the Petri net is associated with the set of p-invariants the place belongs to. This table was generated after the validation and similarity detection of p-invariants, ensuring that no p-invariant shares more than 60% of places in their support.

#### 2. Identification of signaling segments

The p-invariants of *N_GP CR_* are used to create subnets. If the p-invariant support contains more than one place, then a subnet is formed with the places of the support and their connected transitions. One of the subnets of *N_GPCR_*, the subnet *N_GP CR_* for the p-invariant BAR, is shown in Figure 14. A t-invariant analysis is performed on each of the 13 subnets of *N_GPCR_* (four p-invariants of *N_GPCR_* have only one place, thus no subnets are created for them). The 8 t-invariants found in 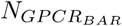 are {*t*_30_*, t*_42_}, {*t*_20_*, t*_42_}, {*t*_3_*, t*_4_}, {*t*_31_*, t*_32_}, {*t*_26_*, t*_25_}, {*t*_13_*, t*_14_}, {*t*_0_*, t*_13_} and {*t*_0_*, t*_4_*, t*_31_*, t*_25_}. The merging of all the t-invariants of 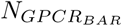having at least one transition in common produces two signaling segments: 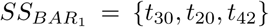 and 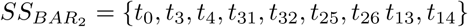. The complete list of signaling segments for the 13 subnets of *N_GP CR_* is given in Table 12.

**Figure 14:**
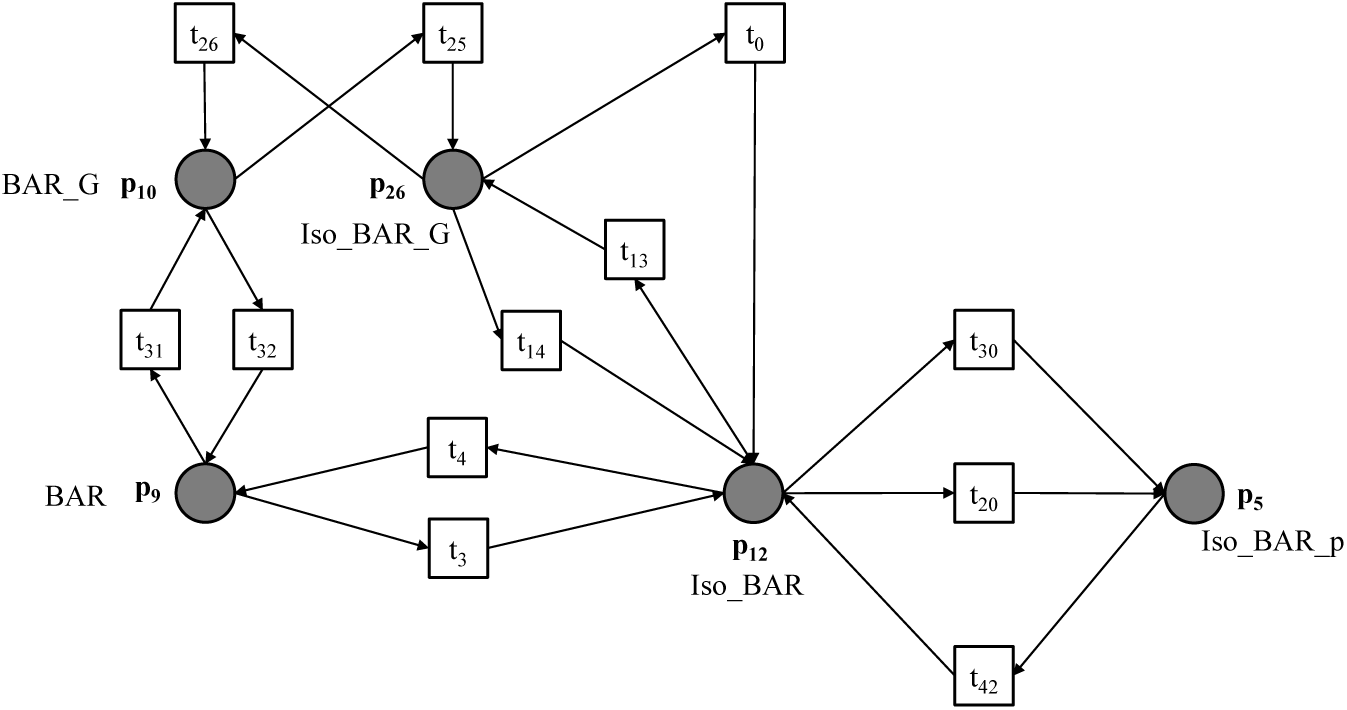
Subgraph 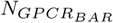 signaling segments. induced by the p-invariant BAR of *N_GPCR_*. This subnet has 8 t-invariants and 2

**Table 12:** The signaling segments for the p-invariants of *N_GPCR_* with more than one place in their support.

| P-invariant | Signaling segments |
| --- | --- |
| Iso | $\{t_{14}, t_{13}, t_0, t_{25}, t_{26}, t_4, t_3\}, \{t_{20}, t_{42}, t_{30}\}$ |
| BAR | $\{t_{20}, t_{42}, t_{30}\}, \{t_{14}, t_4, t_{13}, t_{32}, t_{31}, t_3, t_0, t_{25}, t_{26}\}$ |
| Gbg | $\{t_6, t_7\}, \{t_{14}, t_{13}, t_{32}, t_{31}, t_0, t_{25}, t_{12}, t_{26}\}$ |
| GRK | $\{t_6, t_7\}$ |
| Gs | $\{t_{17}, t_{18}\}, \{t_{14}, t_{13}, t_{32}, t_{11}, t_{31}, t_0, t_{25}, t_{12}, t_{26}\}$ |
| AC | $\{t_{17}, t_{18}\}$ |
| cAMP | $\{t_{37}, t_{36}\}, \{t_{40}, t_{41}\}, \{t_{34}, t_{35}\}, \{t_{39}, t_{38}\}$ |
| PKA | $\{t_{37}, t_{36}\}, \{t_{40}, t_{41}\}, \{t_{39}, t_{38}\}, \{t_{34}, t_{35}\}$ |
| PDE | $\{t_{21}, t_{28}\}$ |
| BRAF | $\{t_9, t_2\}$ |
| MEK | $\{t_{33}, t_{22}\}$ |
| MAPK | $\{t_{27}, t_{24}, t_8, t_{23}\}$ |
| PTP | $\{t_{16}, t_{29}\}$ |

#### 3. Exploration of the Petri net model

The Petri net *N_GP CR_* is explored from place to transition to place using a depth-first search algorithm starting at the place *p*_11_, the signaling source of the model. Crossing a transition is a step. Quadruplets are stored in the table *RFAP*. The first quadruplets from this exploration are [*p*_11_*, p*_12_*, t*_3_, 0] and [*p*_11_*, p*_26_*, t*_25_, 0]. At the following step, the quadruplets are [*p*_12_*, p*_11_*, t*_4_, 1], [*p*_12_*, p*_9_*, t*_4_, 1], [*p*_12_*, p*_5_*, t*_20_, 1], [*p*_12_*, p*_5_*, t*_30_, 1], [*p*_12_*, p*_26_*, t*_13_, 1], [*p*_26_*, p*_11_*, t*_26_, 1], [*p*_26_*, p*_2_*, t*_14_, 1], [*p*_26_*, p*_12_*, t*_0_, 1], [*p*_26_*, p*_12_*, t*_14_, 1], [*p*_26_*, p*_10_*, t*_26_, 1], [*p*_26_*, p*_17_*, t*_0_, 1],and [*p*_26_*, p*_3_*, t*_0_, 1]. At each step, more quadruplets are added to the table *RFAP*. The exploration of *N_GP CR_* is completed at step 7 with the last quadruplets [*p*_15_*, p*_13_*, t*_8_, 7], [*p*_6_*, p*_20_*, t*_5_, 7] and [*p*_24_*, p*_23_*, t*_24_, 7].

### 4.2. Phase II: Creation of the influence graph for the Petri net model of the GPCR-PKA-MAPK signaling network

The three transformation rules of the second phase of the algorithm are executed on the Petri net *N_GPCR_*: the creation of the intermediate interaction graph, the removal of some interaction edges following the signaling segments rule, and the building of the influence graph. The rules use the data structures generated in the first phase.

#### 4. Creation of the interaction graph

First, a vertex is added to the interaction graph for each of the 17 different sets of p-invariants in the lookup table *E_p_*. Then, an interaction edge is added to the graph for each place in the vertices’ supports. For example, the support of the vertex associated with the p-invariant set {AC, Gs} is the place *p*_0_. The values associated with the place *p*_0_ in the lookup table *RFAP* are the quadruplets [*p*_0_*, p*_19_*, t*_18_, 3], [*p*_0_*, p*_3_*, t*_18_, 3] and [*p*_0_*, p*_28_*, t*_10_, 3]. The target places of these quadruplets (*p*_19_, *p*_3_ and *p*_28_) are in the support of the vertices associated to the p-invariant sets {AC}, {Gs} and {cAMP} respectively. Consequently, three edges are added to the graph: {AC, Gs}→{AC}, {AC, Gs}→{Gs}, and {AC, Gs}→{cAMP}. The quadruplet at the origin of an edge is associated with it. In total, 43 edges are added to the interaction graph of *N_GP CR_* and as many entries are made to the lookup table *OE* associating the 8 steps necessary to explore *N_GP CR_* with the quadruplets of the new edges. If a vertex is disconnected from the main graph after every support place have been processed, this vertex is removed. Figure S1 shows the interaction graph as it is created at this point. Table S2 presents the list of the edges with their source and target vertices and the quadruplet associated with each edge. When there is more than one quadruplet in the list, this means that there are as many edges between the two vertices.

#### 5. Removal of interaction edges with the signaling segments rule

Using the lookup table *OE*, exploration steps are iterated from 0 to 7 and the associated quadruplets are tested. At step 0, two edges are processed. The edge from the vertex {Iso} to the vertex {BAR, Iso} with the quadruplet [*p*_11_*, p*_12_*, t*_3_, 0] has for implicit source the p-invariant Iso and for its implicit target the p-invariant BAR. The signaling segment of the p-invariant Iso with the transition *t*_3_ is {*t*_14_*, t*_13_*, t*_0_*, t*_25_*, t*_26_*, t*_4_*, t*_3_}. In the record associated with this signaling segment, the p-invariant {BAR} is added at step 0, and so that recorder is updated. The second edge that is processed is from the vertex {Iso} to the vertex {BAR, Gs, Iso, Gbg} with the quadruplet [*p*_11_*, p*_26_*, t*_25_, 0]. It has for implicit source Iso and for implicit targets the p-invariants BAR, Gs and Gbg. The transition *t*_25_ is in the same signaling segment. BAR is already present in its associated record, but Gs and Gbg are added at step 0, and the recorder is updated. The presence of these p-invariants in the record of the recorder of Iso causes the removal of several other edges associated with later steps. For example, the edges {BAR, Iso} → {Iso} and {BAR, Iso} → {BAR} are removed because their associated quadruplet contains the transition *t*_4_, which is in the previously identified signaling segment. Another example is the edge from the vertex {cAMP, PKA} to the vertex {PKA} with the quadruplet [*p*_34_*, p*_35_*, t*_41_, 5]. Its implicit sources are the p-invariants cAMP and PKA and its implicit target is PKA. However, PKA is already in the record of the signaling segment {*t*_40_, *t*_41_} of cAMP since it was added at step 4 with the edge {cAMP}→ {cAMP, PKA}. Consequently, this edge is also removed.

The quadruplets in red in Table S2 are the edges that are removed by the signaling segments rule. 21 edges remain in the interaction graph representing the Petri net *N_GP CR_*. Figure S2 shows the final interaction graph after the application of the rule.

#### 6. Building the influence graph from the interaction graph

The influence graph of the Petri net *N_GP CR_* is built by processing the interaction graph built at the previous rule. This is accomplished one exploration step at a time with subgraphs of the interaction graph for each exploration step. We present the example of exploration steps 0, 3 and 4. At exploration step 0, the subgraph contains three vertices and the two edges {Iso}→{BAR, Iso} and {Iso}→{BAR, Gs, Iso, Gbg}. This is a maximal bipartite component with a first partition of one source vertex and a second partition with two target vertices. Iso is the core source p-invariant of this component. There is no core target p-invariant. The core source p-invariant is removed from the two target sets. The target is BAR since it is the target set with the lowest cardinality in the second partition. An influence arc is created from Iso to BAR with the two quadruplets [*p*_11_*, p*_12_*, t*_3_, 0] and [*p*_11_*, p*_26_*, t*_25_, 0] and the arc is added to the new influence graph. A condensing relationship is created with Iso being a codomain of BAR. At exploration step 3, the subgraph contains four vertices and the two edges {GRK, Gbg}→{BAR, Iso} and {AC, Gs}→{cAMP}, which already forms two maximal bipartite components. For the first component {GRK, Gbg}→{BAR, Iso}, a previously identified signal condensation relationship causes the p-invariants Gbg and Iso to be removed from the vertices of this component because they are codomains of GRK and BAR, respectively. With one p-invariant left in the source and target partitions each, these p-invariants are the source and target of the new influence arc (GRK→BAR) associated with the quadruplet [*p*_4_*, p*_5_*, t*_30_, 3]. No new condensation relationship is established since this is an enzymatic reaction. For the second component {AC, Gs}→{cAMP}, Gs is removed from the source vertex because of a signal condensation. This p-invariant is a codomain of AC. With one p-invariant left in the source and target partitions each, these p-invariants are the source and target of a new influence arc (AC→cAMP) that is associated with the quadruplet [*p*_0_*, p*_28_*, t*_10_, 3]. No new condensation relationship is established since this is also an enzymatic reaction. At step exploration 4, two new influence arcs are created (GRK→BAR and AC→cAMP), but since there are already arcs connecting these vertices in the influence graph the quadruplets [*p*_21_*, p*_5_*, t*_20_, 4] and [*p*_19_*, p*_28_*, t*_19_, 4] of the new arcs are merged with the quadruplets of the existing arcs. Table 13 shows the correspondence between the interaction graph and the influence graph. Figure S3 represents the unannotated influence graph of the Petri net model *N_GPCR_*.

**Table 13:** This table shows the result of the transformation from the interaction graph to the influence graph representation of *N_GPCR_*. In the interaction graph, each quadruplet represents a different edge. In the influence graph, every quadruplets between the same source and target vertices are associated with a single arc. For example, the arc between Iso and BAR in the influence graph combines two edges from the interaction graph: {Iso} *→* {BAR, Iso} and {Iso} *→* {BAR, Gs, Iso, Gbg}. The quadruplets marked with * are not included in the influence graph because their transitions *t*_13_ and *t*_14_ form a trivial t-invariant. The quadruplet [*p*_26_*, p*_12_*, t*_0_, 1] marked with *†* is not included in the influence graph because its target place *p*_12_ is not in the support of the target vertices Gbg or Gs.

| $v_{source}$ in $G_{interaction}$ | $v_{target}$ in $G_{interaction}$ | Quadruplets | $v_{source}$ in $G_{influence}$ | $v_{target}$ in $G_{influence}$ |
| --- | --- | --- | --- | --- |
| {Iso} | {BAR, Iso} | $[p_{11}, p_{12}, t_3, 0]$ | Iso | BAR |
| {Iso} | {BAR, Gs, Iso, Gbg} | $[p_{11}, p_{26}, t_{25}, 0]$ | | |
| {BAR, Iso} | {BAR, Gs, Iso, Gbg} | $[p_{12}, p_{26}, t_{13}, 1]^*$ | BAR | Gbg |
| {BAR, Gs, Iso, Gbg} | {Gbg} | $[p_{26}, p_{17}, t_0, 1]$ | | |
| {BAR, Gs, Iso, Gbg} | {Gs, Gbg} | $[p_{26}, p_2, t_{14}, 1]^*$ | | |
| {BAR, Gs, Iso, Gbg} | {BAR, Iso} | $[p_{26}, p_{12}, t_{14}, 1]^*, [p_{26}, p_{12}, t_0, 1]^\dagger$ | | |
| {BAR, Gs, Iso, Gbg} | {Gs} | $[p_{26}, p_3, t_0, 1]$ | BAR | Gs |
| {BAR, Iso} | {BAR, Gs, Iso, Gbg} | $[p_{12}, p_{26}, t_{13}, 1]^*$ | | |
| {BAR, Gs, Iso, Gbg} | {Gs, Gbg} | $[p_{26}, p_2, t_{14}, 1]^*$ | | |
| {BAR, Gs, Iso, Gbg} | {BAR, Iso} | $[p_{26}, p_{12}, t_{14}, 1]^*, [p_{26}, p_{12}, t_0, 1]^\dagger$ | | |
| {Gs} | {AC, Gs} | $[p_3, p_0, t_{17}, 2]$ | Gs | AC |
| {Gbg} | {GRK, Gbg} | $[p_{17}, p_4, t_6, 2]$ | Gbg | GRK |
| {AC} | {cAMP} | $[p_{19}, p_{28}, t_{19}, 4]$ | AC | cAMP |
| {AC, Gs} | {cAMP} | $[p_0, p_{28}, t_{10}, 3]$ | | |
| {GRK, Gbg} | {BAR, Iso} | $[p_4, p_5, t_{30}, 3]$ | GRK | BAR |
| {GRK} | {BAR, Iso} | $[p_{21}, p_5, t_{20}, 4]$ | | |
| {cAMP} | {cAMP, PKA} | $[p_{28}, p_{34}, t_{40}, 4], [p_{28}, p_{31}, t_{34}, 4], [p_{28}, p_{32}, t_{36}, 4], [p_{28}, p_{33}, t_{38}, 4]$ | cAMP | PKA |
| {cAMP, PKA} | {BRaf} | $[p_{31}, p_{16}, t_9, 5]$ | PKA | Braf |
| {cAMP, PKA} | {PTP} | $[p_{31}, p_{25}, t_{16}, 5]$ | PKA | PTP |
| {cAMP, PKA} | {PDE} | $[p_{31}, p_{30}, t_{21}, 5]$ | PKA | PDE |
| {PTP} | {MAPK} | $[p_{25}, p_{23}, t_{23}, 6], [p_{24}, p_{23}, t_{24}, 7]$ | PTP | MAPK |
| {BRaf} | {MEK} | $[p_{16}, p_{15}, t_{22}, 6]$ | BRaf | MEK |
| {PDE} | {cAMP} | $[p_{30}, p_{20}, t_1, 6], [p_6, p_{20}, t_5, 7]$ | PDE | cAMP |
| {MEK} | {MAPK} | $[p_{15}, p_{13}, t_8, 7]$ | MEK | MAPK |

### 4.3. Phase III: Annotation of the influence graph for the Petri net model of the GPCR-PKA-MAPK signaling network

The three rules of the final phase of the algorithm are executed for the Petri net *N_GP CR_* and annotate the influence graph created at the second phase: detecting the signaling places for each vertex of the influence graph, associating transitions from the Petri net model with the arcs of the influence graph, and assigning a type (activation or inhibition) to the arcs.

#### 7. Detecting signaling places

The annotation of the influence graph starts by detecting the signaling places of the different vertices. A signaling place corresponds to an active state of the molecule. This rule first identifies the leaves of the influence graph. In the influence graph of the Petri net *N_GP CR_*, the vertex MAPK is the only leaf. Since this vertex is connected to the vertices MEK and PTP by enzymatic reactions, no verification of the graph is necessary. The algorithm continues by identifying the signaling places of each vertex by applying the criteria of Definition 19 to the quadruplets of the arcs starting from that vertex. For the vertex Gs, the processing of the quadruplet [*p*_3_*, p*_0_*, t*_17_, 2] of the arc to the vertex AC detects *p*_3_ and *p*_0_ as signaling places for Gs because both places are in its support. For the vertex cAMP, the processing of the quadruplets [*p*_28_*, p*_34_*, t*_40_, 4], [*p*_28_*, p*_31_*, t*_34_, 4], [*p*_28_*, p*_32_*, t*_36_, 4] and [*p*_28_*, p*_33_*, t*_38_, 4] of the arc to the vertex PKA detects *p*_28_, *p*_34_, *p*_31_, *p*_32_ and *p*_33_ as signaling places of this vertex because all places are in the support of cAMP. For the vertex MEK, the processing of the quadruplet [*p*_15_*, p*_13_*, t*_8_, 7] of the arc to the vertex MAPK detects *p*_15_ as the signaling place of this vertex. The target place *p*_13_ is not labeled as a signaling place because it is not in the support of the vertex MEK. For the leaf vertex MAPK, the user is prompted to identify its signaling places since it has no outgoing arc. The place *p*_13_ is selected. The next test is to search for an unbinding reaction caused by the signal. The transition *t*_0_ has one preplace and three postplaces. It is associated with arcs from the vertex BAR. With the criteria of Definition 19, only the place *p*_26_ is identified as a signaling places. However, it is possible that *p*_12_ can also be considered an active state for the receptor since it is bound to the agonist Iso. Because of this undecidability, the user is prompted and *p*_12_ is added to the list of signaling places of the vertex BAR. After this processing, four vertices have a list of signaling places that includes their entire support: GRK, AC, PTP and PDE. This situation is caused by a modeling choice from the authors of the model: the inactive state of these enzymes is modeled to have a basal level of activity instead of having absolutely no effect at all. For the cyclase AC, this is shown by the transition *t*_19_ which represents the basal catalytic conversion of ATP into cAMP. The user is prompted and only *p*_0_ is labeled as a signaling place of this vertex. To complete the first annotation rule of the influence graph of the Petri net *N_GP CR_*, all the places of the signal vertex Iso are labeled as signaling places. Also, the places that were not detected as signaling places are labeled as non-signaling places. The signaling places for the vertices of the influence graph are listed in Table 14.

**Table 14:**
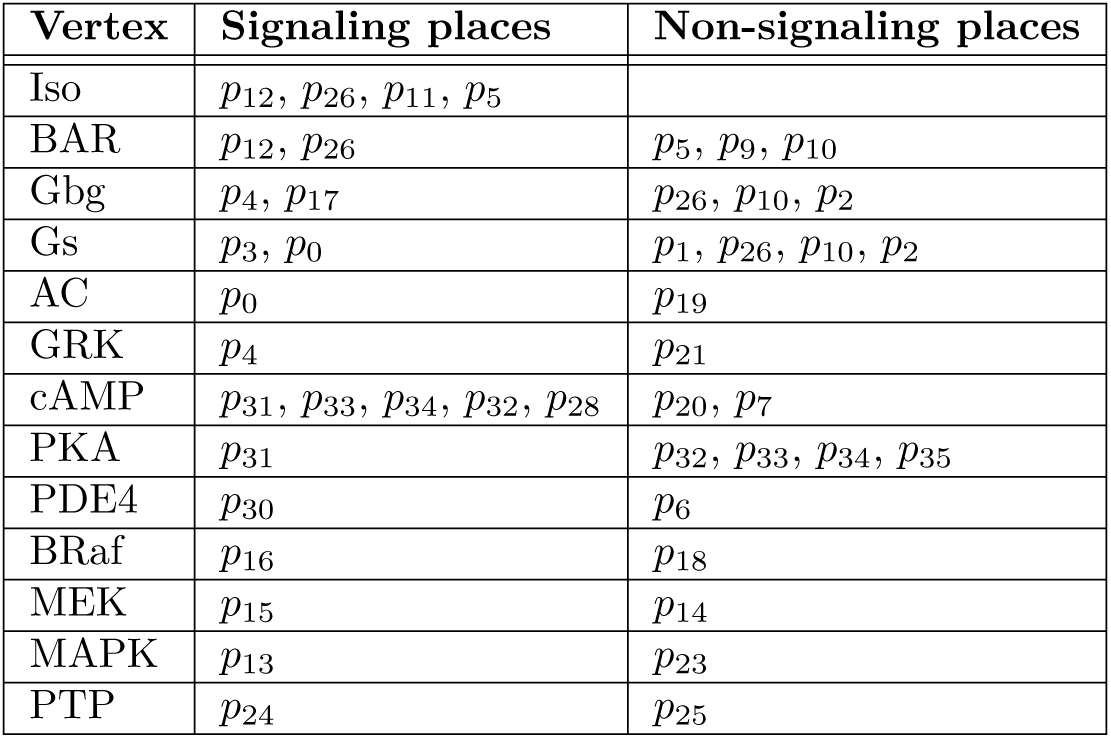
The signaling and the non-signaling places of the vertex of the influence graph of the Petri net *N_GPCR_* as identified by the rule 3.3.1.

#### 8. Associating signaling transitions to influence arcs

After annotating the influence graph vertices with signaling places, its arcs are annotated with the transitions transmitting the signal in the model. Each arc of the influence graph of the Petri net *N_GP CR_* is tested with the conditions of Definition 20. Four quadruplets are removed. For the arc AC→cAMP, the quadruplet with the transition *t*_19_ is removed because the source place of the quadruplet is a non-signaling place of the vertex AC. For the same reason, the quadruplets with the transitions *t*_5_, *t*_23_ and *t*_20_ are removed from the arcs PDE→cAMP, PTP→MAPK and GRK→BAR, respectively. In total, 19 quadruplets remain in the influence graph and the signaling transitions are associated with the arcs.

#### 9. Determining the type of influence arcs

In the last rule of the transformation algorithm, the influence type of the arcs of the graph is determined with Definition 21. Of the 15 influence arcs *e*(*v_source_, v_target_*) of the graph representation of the Petri net *N_GP CR_*, four arcs respect the condition of having a quadruplet [*p_source_, p_target_, t, step*] with *p_target_* being a non-signaling place of *v_target_* and all preplaces of *t* other than *p_source_* are signaling places of *v_source_*. The influence type of these arcs is inhibitory.

These four arcs are: 1) GRK→BAR where *p*_5_ is a non-signaling place of the vertex BAR and *p*_12_ is a signaling place of the same vertex for the quadruplet [*p*_4_*, p*_5_*, t*_30_, 3], 2) PDE→cAMP where *p*_20_ is a non-signaling place of the vertex cAMP and *p*_28_ is a signaling place of the same vertex for the quadruplet [*p*_30_*, p*_20_*, t*_1_, 6], 3) PKA→PTP where *p*_25_ is a non-signaling place of the vertex PTP and *p*_25_ is a signaling place of the same vertex for the quadruplet [*p*_31_*, p*_25_*, t*_16_, 5] and 4) PTP→MAPK where *p*_23_ is a non-signaling place of the vertex MAPK and *p*_13_ is a signaling place of the same vertex for the quadruplet [*p*_24_*, p*_23_*, t*_24_, 7]. The other arcs of the graph represent an activation influence. For example, the arc BAR→Gs with the quadruplet [*p*_26_*, p*_3_*, t*_0_, 1] has a target place *p*_3_ that is a signaling place, thus the arc has an activation influence. The final influence graph of the Petri net *N_GP CR_* is shown in Figure 15.

**Figure 15:**
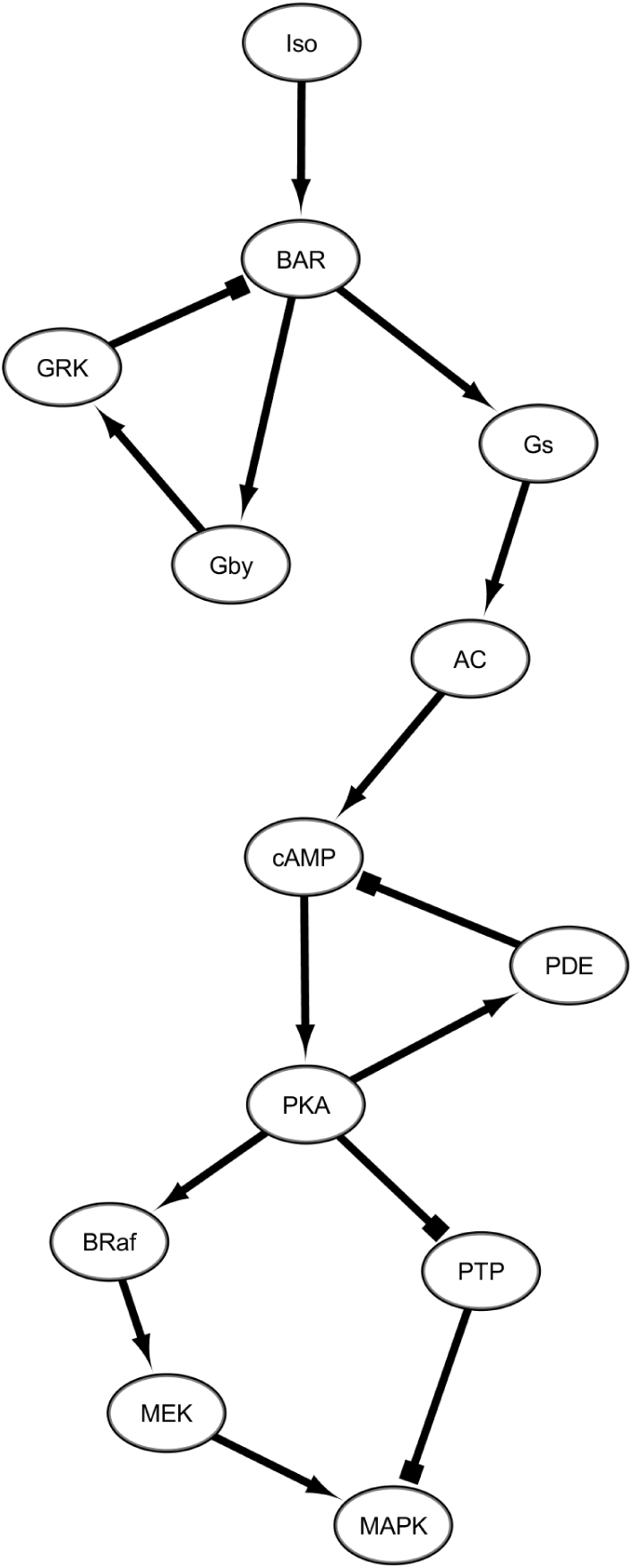
The annotated influence graph of the Petri net *N_GPCR_* created by the transformation algorithm. Activation arcs are represented by arrows and inhibition arcs have T-shapped heads.

## 5. Concluding remarks

The algorithm presented in this paper can be used to transform a biological model represented as a Petri net into an annotated influence graph. The resulting graph is useful to study and validate the topology of the model’s network. This representation is also particularly useful for signaling systems to understand how the regulatory network motifs and their interconnections process biological signals. With the annotations, it becomes possible to map simulation data onto the influence graph to create a dynamic representation of the simulated model, or what we call a dynamic graph (find examples in [35] and [36]). Following this idea, if the model is a continuous or hybrid Petri net with kinetic information and initial conditions that can be simulated, then the resulting time series can be used to color the graph vertices and to modify the width of the arcs in a time-dependent manner as molecules are activated and deactivated. The state variables of the model are the signaling places of the graph, and the reaction rates are the signaling transitions. The output of this transformation is a systemic perspective of a biological dynamical model and one can see how the dynamics of regulatory motifs such as desensitization, crosstalk, switch-like behavior, overshoot correction, frequency detection or noise filtering can lead to a complex biological response [30]. The transformation example presented in Section 4 is a Petri net model of 34 places and 38 transitions. After the transformation, the resulting influence graph, with 13 vertices and 15 arcs, represents the underlying biological entities and reactions of the model. The two negative feedback loops and the coherent feedforward motif of this signaling pathway become obvious. By combining simulation data from this model with the influence graph, it becomes possible to observe when the loops are active and understand how the signal is processed by this pathway. Every mathematical model of a signaling network respecting the input constraints of the algorithm can be transformed into an influence graph by first converting it to a Petri net model, with a tool such as Snoopy, which has a SBML import feature [42].

This first version of the transformation algorithm has some limitations. Despite the fact that many signals might be simultaneously stimulating a biological system, dealing with only one source of signal is a limitation of the algorithm in its current form. Also, enzymatic reactions must be represented by a single Michaelis-Menten reaction, not by three mass action reactions. In the Petri net, this means that there is only one transition per enzymatic reaction and a read arc linking the enzyme to the transition. To allow for the transformation of models using complete enzymatic representations, a new transformation rule could automatically convert these three-step reactions into a single one. Dealing with larger complexes is also problematic for the current algorithm, particularly if allosteric modulators regulate the activity of enzymes. The direction of these binding interactions is tricky to interpret. Maximal bipartite decomposition might be useful to identify these reactions in a model, and this will be considered in future developments of the algorithm. When information is missing, the current algorithm sends requests to a user. This information is sometimes present in the kinetic parameters of the mathematical model. Consequently, another possible improvement of the algorithm is to explore if a Petri net with kinetic parameters, such as a continuous Petri net, could provide this information to the algorithm and thus reduce the need for requests. The algorithm can already transform several biological signaling models and promises to be a practical and handy new tool for modeling in systems biology.

## 7. Declaration of generative AI in scientific writing

The authors did not use any AI or AI-assisted technologies when writing the content of this manuscript.

## 8. Acknowledgments

We are grateful for the useful comments formulated by the anonymous reviewers of this paper. Their observations and suggestions helped us to improve our manuscript. This work was funded by a Discovery grant from the Natural Sciences and Engineering Research Council of Canada awarded to Simon V. Hardy.

## 9. Author contributions statement

SV Hardy proposed the idea. The algorithm was mainly developed by S Gamache-Poirier and A Souvane with later contributions by W Leclerc and C Villeneuve. W Leclerc wrote a first version of the algorithm description and reviewed the manuscript. SV Hardy wrote and reviewed the manuscript.

## Supplementary material

### Detailed description of the construction of the lookup table RFAP from example 4

At the first step of the DFS of the Petri net *N*_1_ from the source place *p*_1_ (step 0), *t*_0_ is crossed and the target place *p*_3_ is reached. The quadruplet [*p*_1_*, p*_3_*, t*_0_, 0] is added in the lookup table *RFAP* for the place *p*_1_. The transition *t*_0_ has no other post-place and *p*_1_ has no other post-transition, thus *p*_1_ is set as visited, and the unvisited target place *p*_3_ becomes the source place of the next step. At the second step of the DFS (step 1), *t*_1_ is crossed from the source place *p*_3_ and the target places *p*_1_ and *p*_2_ are reached. At the same step, *t*_2_ is crossed from the source place *p*_3_ and the target place *p*_5_ is reached. The quadruplets [*p*_3_*, p*_1_*, t*_1_, 1], [*p*_3_*, p*_2_*, t*_1_, 1] and [*p*_3_*, p*_5_*, t*_2_, 1] are added to the lookup table *RFAP* for the place *p*_3_. The transitions *t*_1_ and *t*_2_ have no other post-place and *p*_3_ has no other post-transition, thus *p*_3_ is set as visited, and the unvisited target places *p*_2_ and *p*_5_ become the source places of the next step of the DFS (step 2). From *p*_2_, *t*_0_ is crossed to reach *p*_3_, which adds the quadruplet [*p*_2_*, p*_3_*, t*_0_, 2] to the table *RFAP*. The place *p*_2_ has no other post-transition and is set as visited. From *p*_5_, *t*_3_ is crossed to reach *p*_4_, which adds the quadruplet [*p*_5_*, p*_4_*, t*_3_, 2] to the table *RFAP*. The place *p*_5_ has no other post-transition and is set as visited, and the unvisited target place *p*_4_ becomes the source places of the next step of the DFS (step 3). From *p*_4_, *t*_2_ is crossed to reach *p*_5_, which adds the quadruplet [*p*_4_*, p*_5_*, t*_2_, 3] to the table *RFAP*. The place *p*_4_ has no other post-transition and is set as visited. Since *p*_5_ has already been visited, the DFS ends. Table 3 of the paper is the *RFAP* lookup table for the DFS of the Petri net *N*_1_.

### Detailed description of the removal of edges with the signaling segments rule from example 10

Let’s apply the signaling segments rule to the Petri net *N*_1_ to decide if certain edges of the interaction graph have to be removed following Algorithm 2.1. Considering the lookup table *OE* (Table 5), the first edge in the interaction graph to be considered according to its exploration step is the edge from the vertex *v*_1_ to the vertex *v*_3_ with its associated quadruplet [*p*_1_*, p*_3_*, t*_0_, 0]. In the lookup table *placeT oInvariant*, the source place *p*_1_ of the quadruplet is associated with the p-invariant set {*y*_1_} and the target place *p*_3_ to the p-invariant set {*y*_1_*, y*_2_}. After applying the function implicitConnectedInv to these two sets, the source p-invariant set is not modified because {*y*_1_} ≠ {*y*_1_*, y*_2_} but {*y*_1_} ⊂ {*y*_1_*, y*_2_}, thus it is still {*y*_1_}; but the target p-invariant set is modified to {*y*_2_}, which is the result of {*y*_1_*, y*_2_} − {*y*_1_}, since {*y*_1_*, y*_2_} ≠ {*y*_1_} and {*y*_1_*, y*_2_} ⊄ {*y*_1_}. Then, using invariantToRecorders, the set of recorders for the target *y*_2_ is accessed. This set contains only one recorder because there is one signaling segment ({*t*_0_*, t*_1_}) associated with this p-invariant. Its record is still empty at this point. Since the source p-invariant *y*_1_ is not in this record, the p-invariant *y*_2_ is consequently added at the 0*^th^* step of the record for this signaling segment in the recorder of the source p-invariant. The edge is thus kept in the interaction graph. To summarize, if the signaling segment has not already been used to transduce the signal from the target p-invariant to the source p-invariant, the target p-invariant is registered in the source recorder for the corresponding signaling segment at the n*^th^* step. As intended by this rule, this will prevent this signaling segment to be used at a later step. Since no other edge shares this source place and transition, the algorithm is ready to evaluate another edge. It continues with one of the three edges at exploration step 1. For the edge from *v*_3_ to *v*_1_ with the associated quadruplet [*p*_3_*, p*_1_*, t*_1_, 1], the implicit source p-invariant set is {*y*_2_} and the target p-invariant set is {*y*_1_}. The set of recorders for target p-invariant *y*_1_ is consequently accessed. Since the recorder of the signaling segment {*t*_0_*, t*_1_} now contains the *y*_2_, which is the source p-invariant of this quadruplet, this edge is removed from the interaction graph. Subsequently, because the quadruplet [*p*_3_*, p*_2_*, t*_1_, 1] of the edge from *v*_3_ to *v*_2_ contains the same source place and transition, this edge is also removed. The last edge of step 1, which connects the vertices *v*_3_ and *v*_4_ and is associated with the quadruplet [*p*_3_*, p*_5_*, t*_2_, 1], is kept in the interaction graph because neither of its source p-invariants *y*_1_ and *y*_2_ is in the recorder of the target p-invariant *y*_3_ for the signaling segment {*t*_2_*, t*_3_}. However, in this case, since this signaling segment is not in the recorders of source p-invariants *y*_1_ and *y*_2_, the target p-invariant *y*_3_ is not added at the 1*^st^* step of the recorders. The final edge to be considered connects the vertices *v*_2_ and *v*_3_ and has [*p*_2_*, p*_3_*, t*_0_, 2] as its quadruplet. Its source p-invariant set is {*y*_2_} and its implicit target set is {*y*_1_}. Because the source p-invariant *y*_2_ is already in the recorder associated with *y*_1_, for the signaling segment {*t*_0_*, t*_1_}, this edge is removed from the graph. In summary, after applying the signaling segments rule to the interaction graph of the Petri net *N*_1_, two edges are kept and three edges are removed. The edges with the quadruplets [*p*_3_*, p*_1_*, t*_1_, 1], [*p*_3_*, p*_2_*, t*_1_, 1] and [*p*_2_*, p*_3_*, t*_0_, 2] are all removed because the edge [*p*_1_*, p*_3_*, t*_0_, 0] is first considered and that *t*_0_ and *t*_1_ are part of the same signaling segment. The resulting interaction graph is shown in Figure 7 of the paper. The state of the lookup table invariantToRecorders after the application of the rule is shown in Table 6 of the paper.

#### Algorithm S1 Preparatory phase and Petri net analysis (phase I)

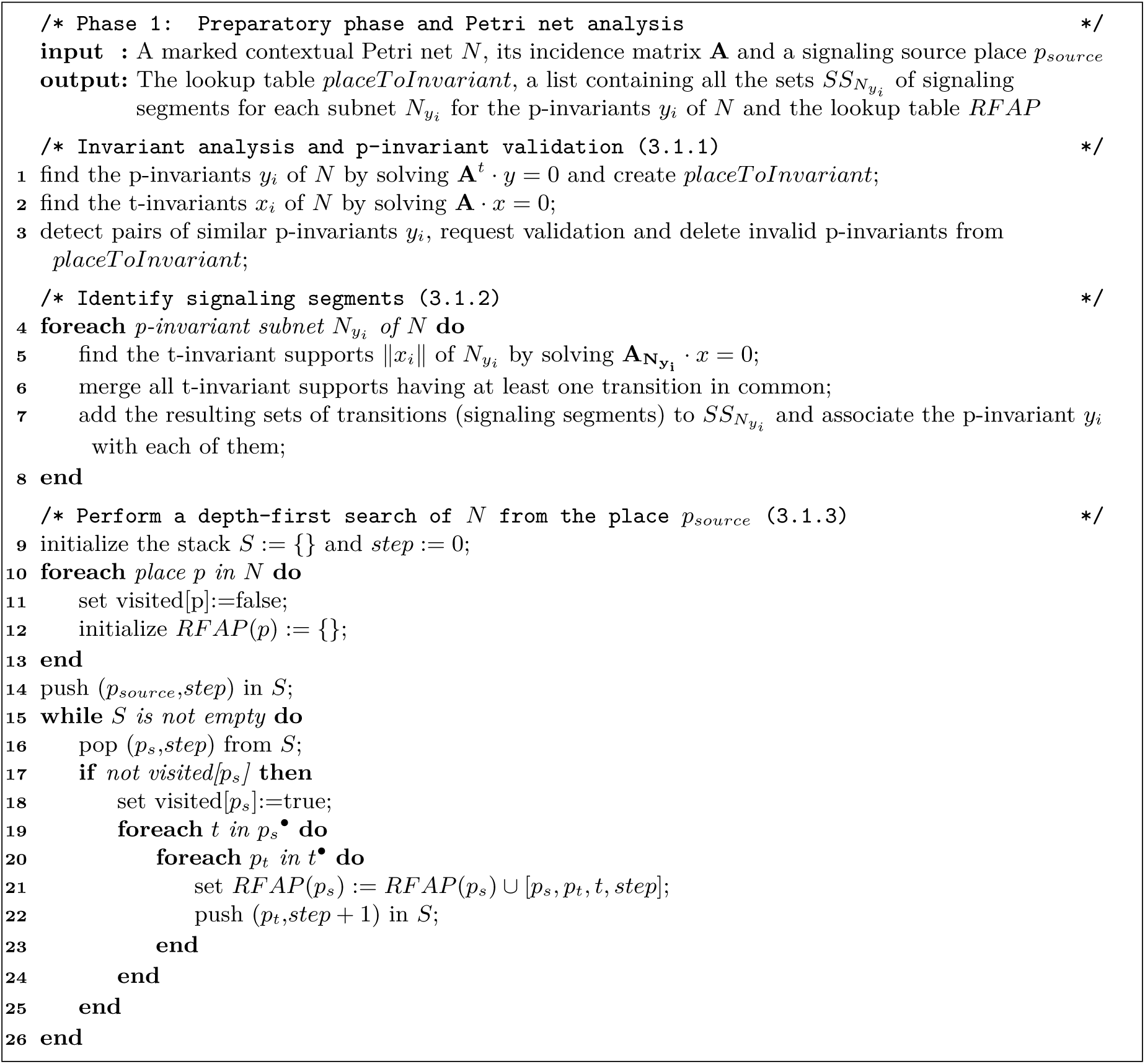

#### Algorithm S2 Building the influence graph from the interaction graph (phase II)

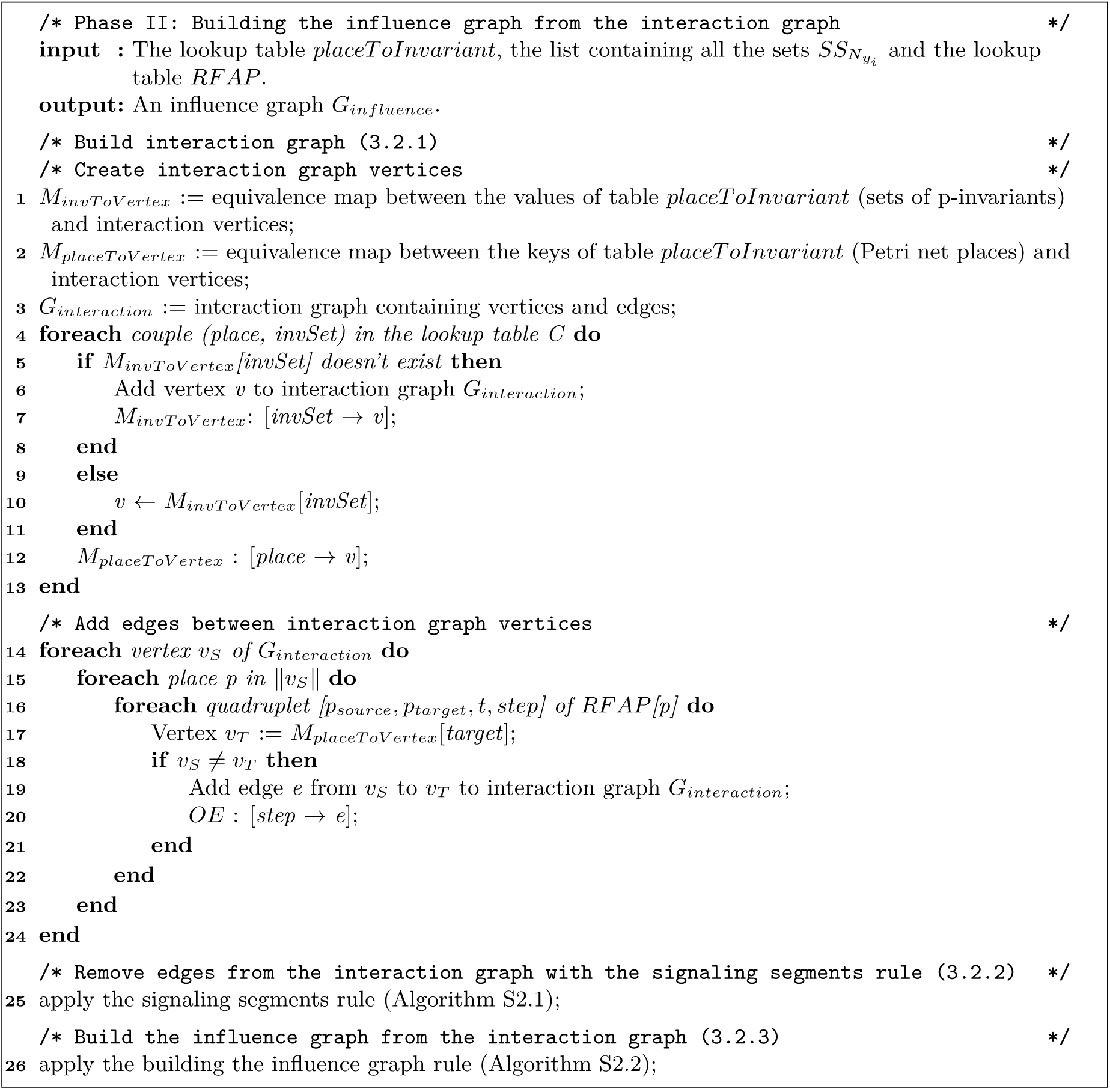

#### Algorithm S2.1 Algorithm for removing interaction edges with the signaling segments rule

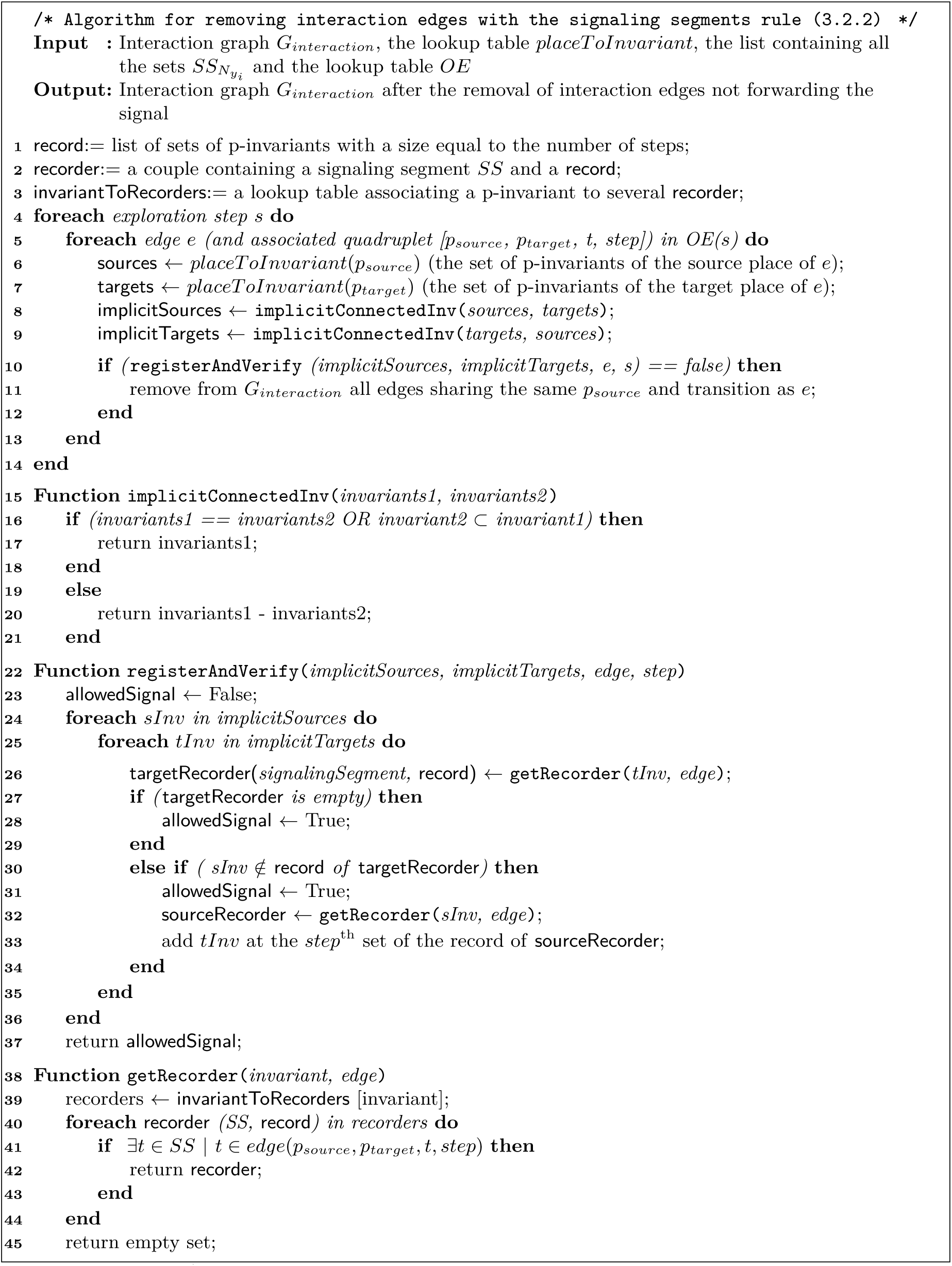

#### Algorithm S2.2 Algorithm for building the influence graph.

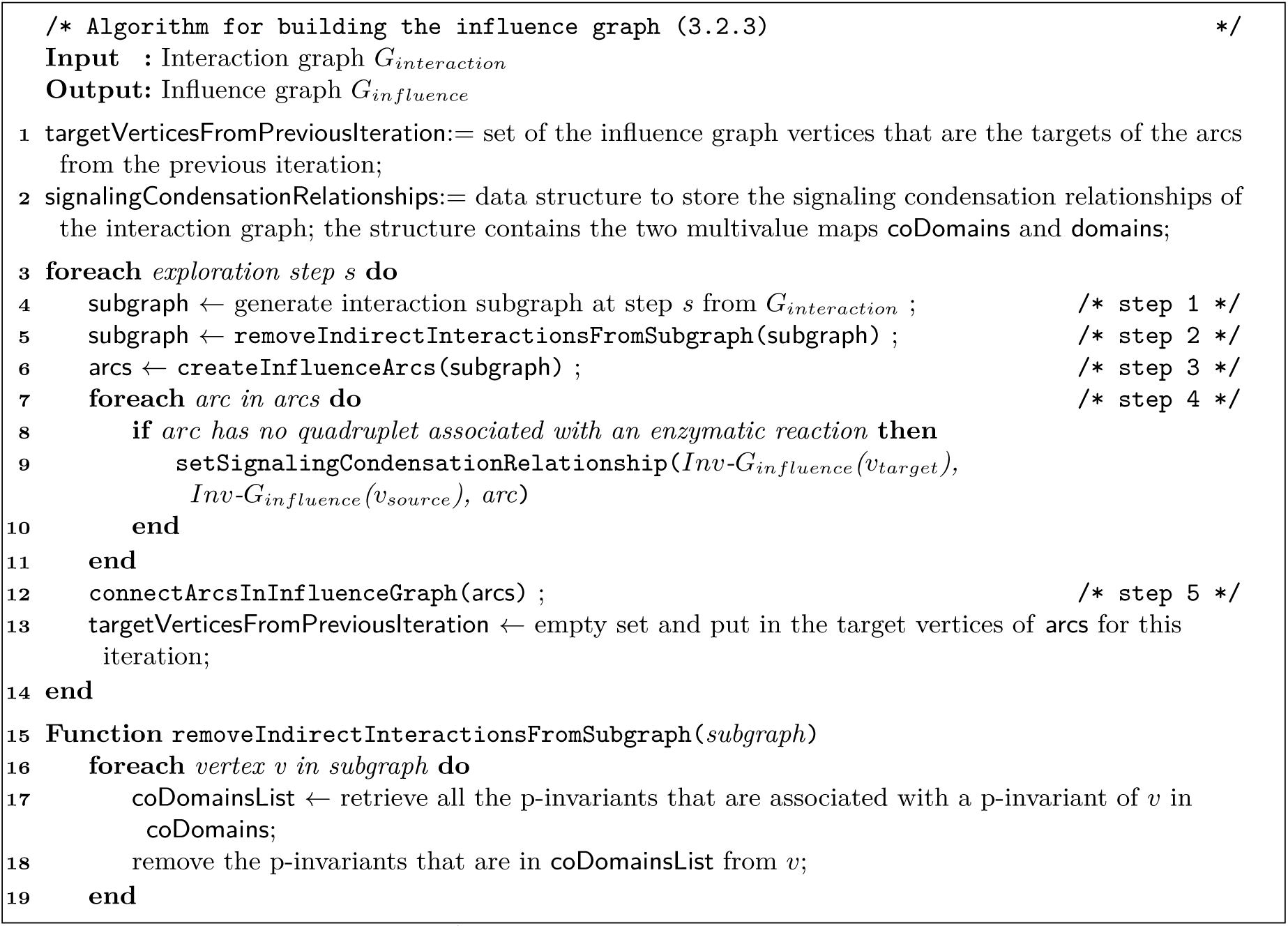
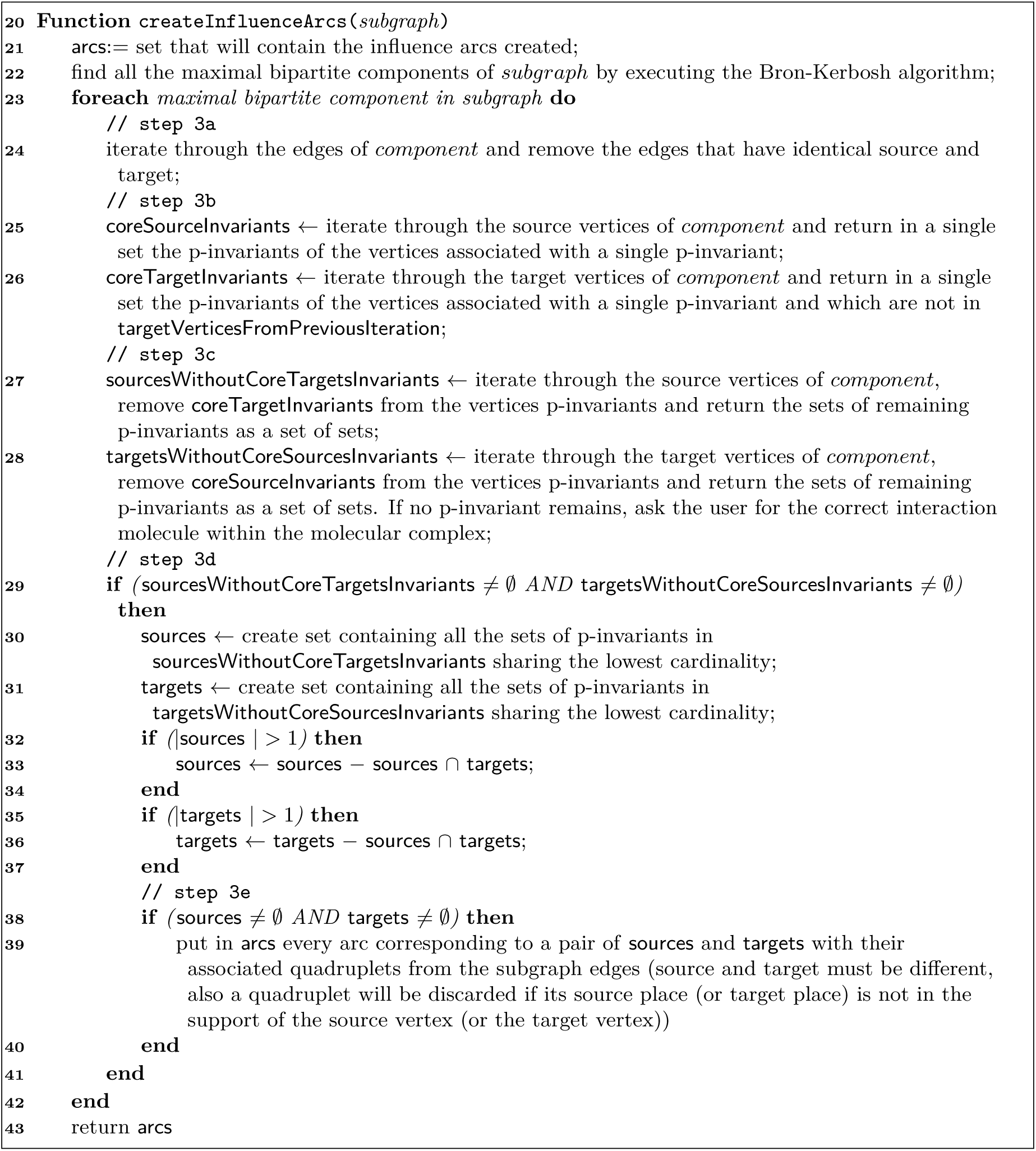
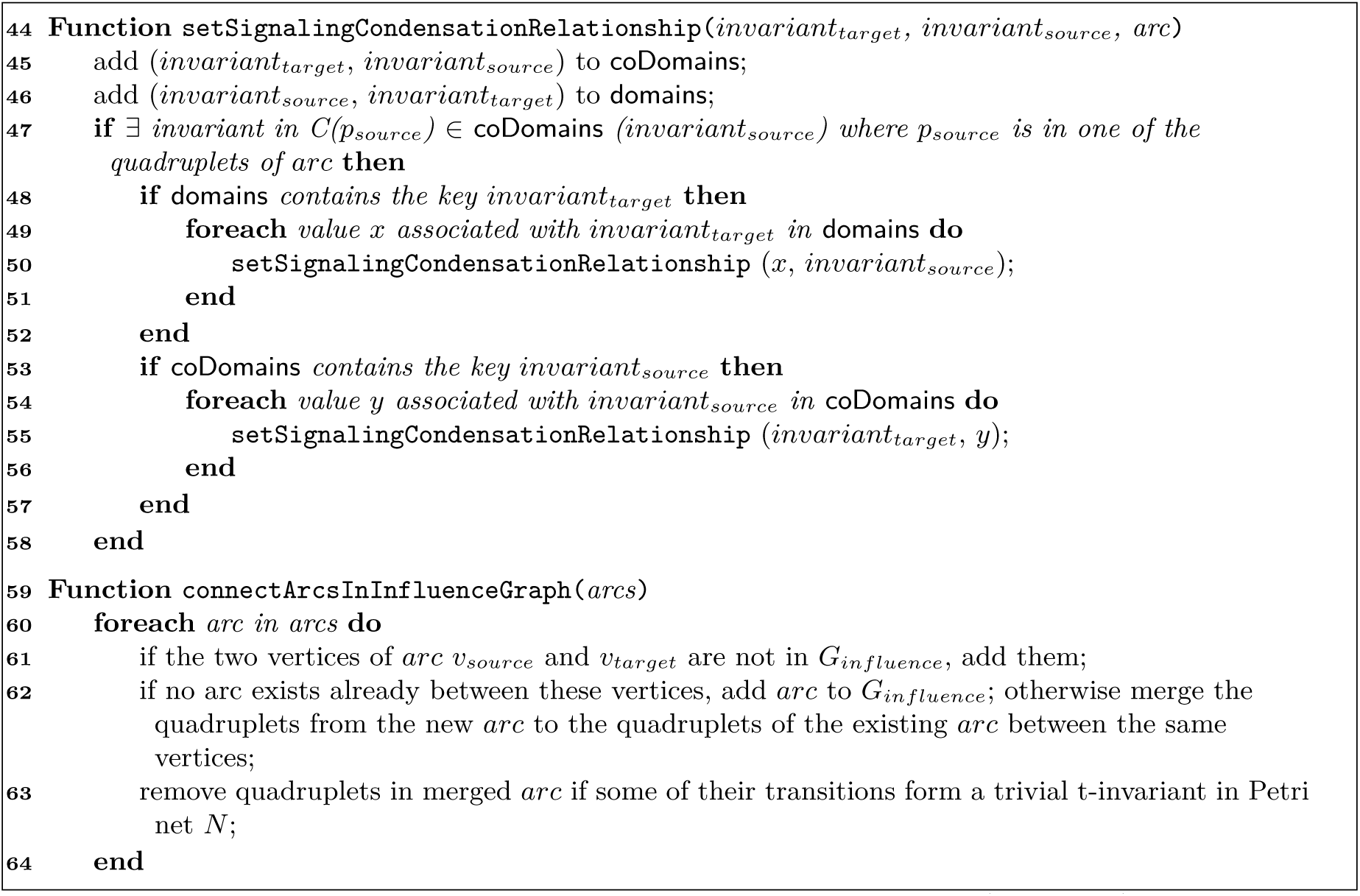

#### Algorithm S3 Annotating the influence graph

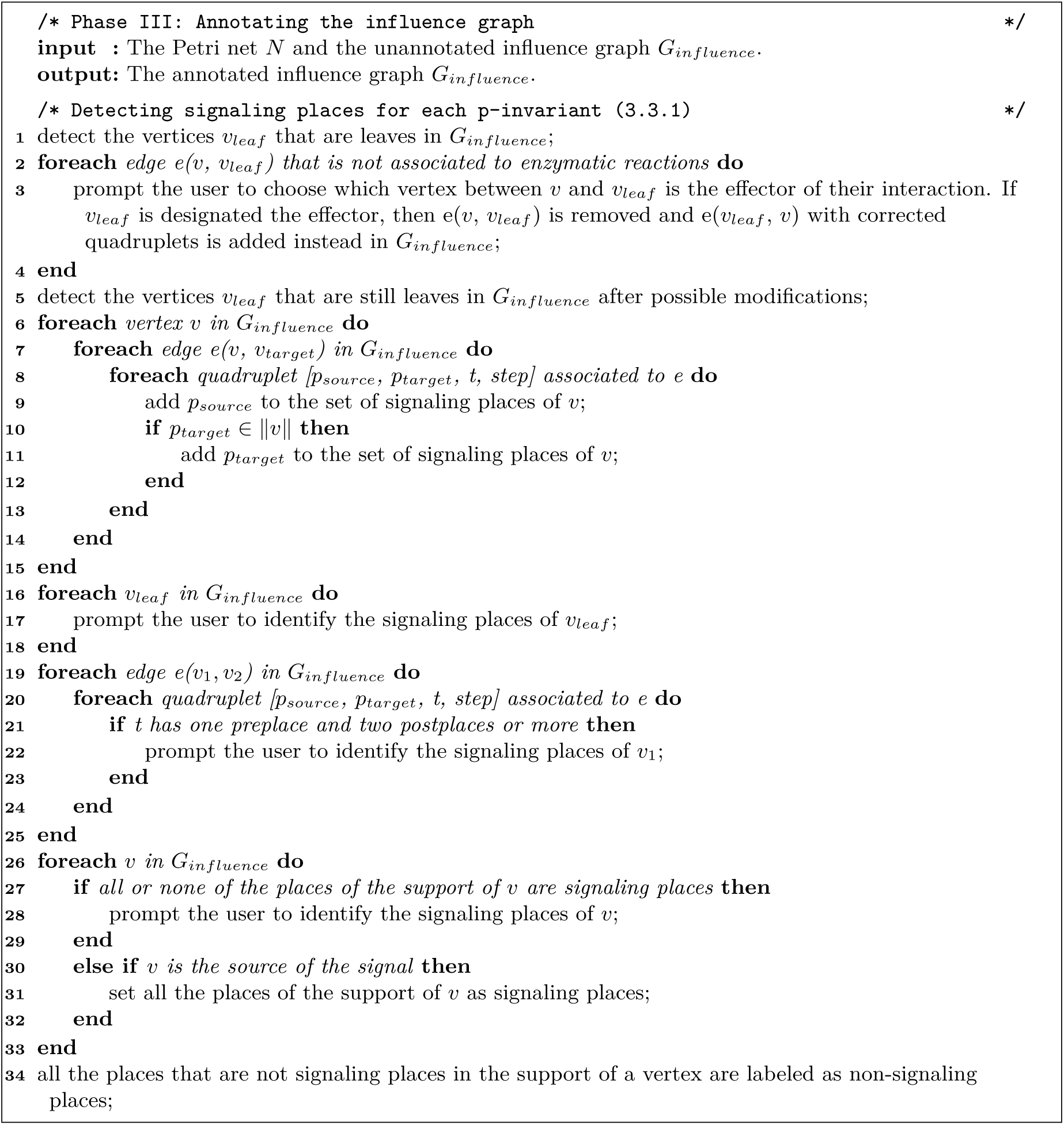
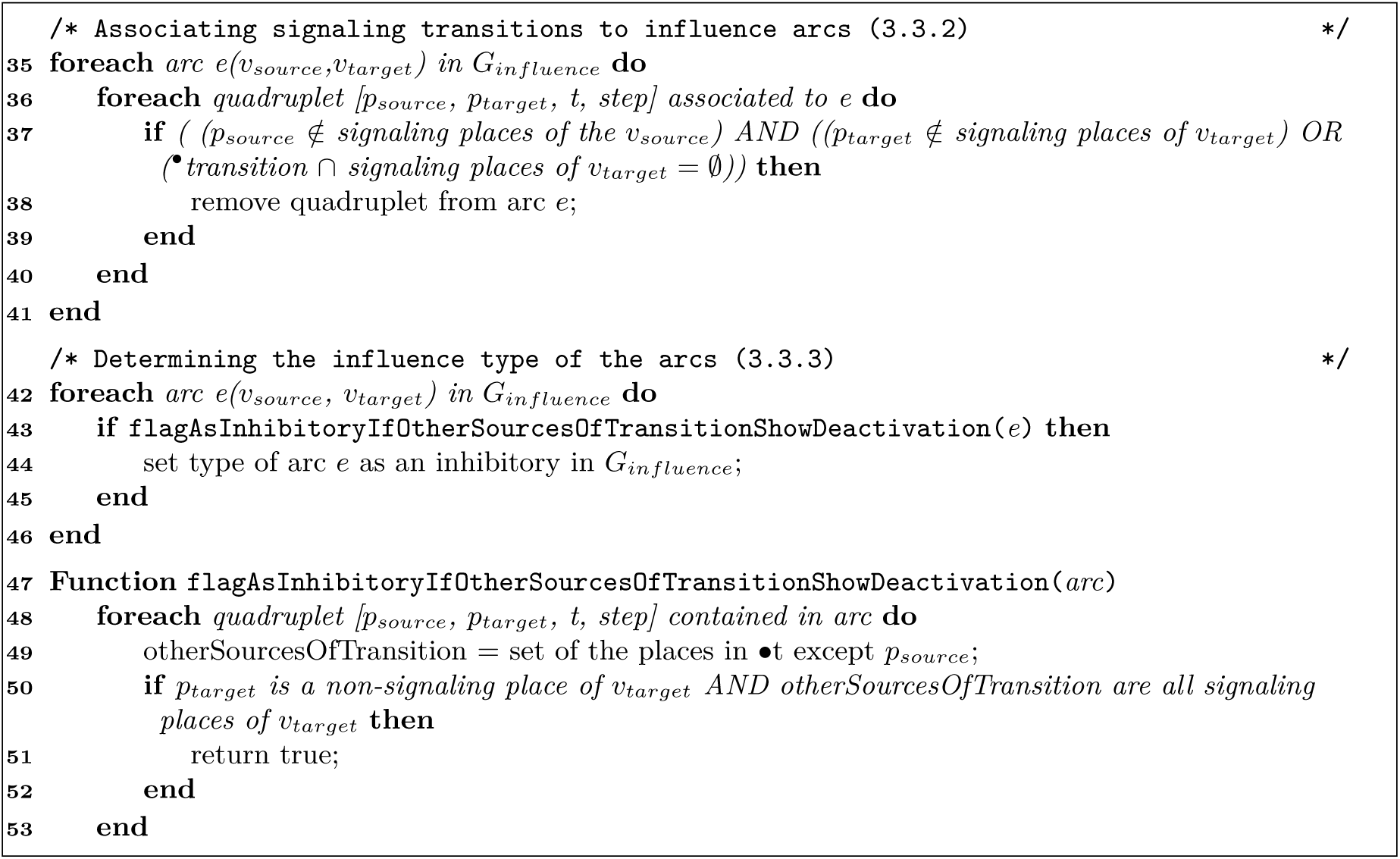

**Table S1:** The lookup table *placeT oInvariant* the Petri net *N_GPCR_* associating each of its places to the set of p-invariants the place belongs to.

| Places | $E_p$, set of p-invariants |
| --- | --- |
| $p_{11}$ : iso_extra | {ISO} |
| $p_5$ : iso_BAR_p_cyto_mem | {ISO, BAR} |
| $p_{12}$ : iso_BAR_cyto_mem | {ISO, BAR} |
| $p_{26}$ : iso_BAR_G_cyto_mem | {ISO, BAR, Gs, Gbg} |
| $p_4$ : GRK_bg_cyto | {Gbg, GRK} |
| $p_{17}$ : bg_cyto | {Gbg} |
| $p_{21}$ : GRK_cyto | {GRK} |
| $p_9$ : BAR_cyto_mem | {BAR} |
| $p_{10}$ : BAR_G_cyto_mem | {BAR, Gs, Gbg} |
| $p_2$ : G_protein_cyto | {Gbg, Gs} |
| $p_3$ : G_a_s_cyto | {Gs} |
| $p_1$ : G_GDP_cyto | {Gs} |
| $p_0$ : AC_active_cyto_mem | {AC, Gs} |
| $p_{19}$ : AC_cyto_mem | {AC} |
| $p_{35}$ : R2C2_cyto | {PKA} |
| $p_{32}$ : c3_R2C2_cyto | {cAMP, PKA} |
| $p_{34}$ : c_R2C2_cyto | {cAMP, PKA} |
| $p_{31}$ : PKA_cyto | {cAMP, PKA} |
| $p_{33}$ : c2_R2C2_cyto | {cAMP, PKA} |
| $p_{20}$ : AMP_cyto | {cAMP} |
| $p_7$ : ATP_cyto | {cAMP} |
| $p_{28}$ : cAMP_cyto | {cAMP} |
| $p_6$ : PDE4_cyto | {PDE} |
| $p_{30}$ : PDE4_P_cyto | {PDE} |
| $p_{25}$ : PTP_PKA_cyto | {PTP} |
| $p_{24}$ : PTP_cyto | {PTP} |
| $p_{13}$ : MAPK_active_cyto | {MAPK} |
| $p_{23}$ : MAPK_cyto | {MAPK} |
| $p_{14}$ : MEK_cyto | {MEK} |
| $p_{15}$ : MEK_active_cyto | {MEK} |
| $p_{16}$ : B_Raf_active_cyto | {Braf} |
| $p_{18}$ : B_Raf_cyto | {Braf} |
| $p_{27}$ : PDE_high_km_cyto | {PDE<sub>highKm</sub>} |
| $p_{29}$ : PTP_PP_cyto | {PP_PTP} |
| $p_8$ : PP_PDE_cyto | {PP_PDE} |
| $p_{22}$ : PP2A_cyto | {PP2A} |

**Table S2:** The edges of the interaction graph of the Petri net *N_GPCR_* with the source vertex, the target vertex and the associated quadruplets. Each quadruplet corresponds to a different edge between the same source and target. In red, the edges that are removed by the signaling segments rule.

| Source vertex | Target vertex | Quadruplets |
| --- | --- | --- |
| {Iso} | {BAR, Iso} | $[p_{11}, p_{12}, t_3, 0]$ |
| {Iso} | {BAR, Gs, Iso, Gbg} | $[p_{11}, p_{26}, t_{25}, 0]$ |
| {BAR, Iso} | {BAR, Gs, Iso, Gbg} | $[p_{12}, p_{26}, t_{13}, 1]$ |
| {BAR, Iso} | {BAR} | $[p_{12}, p_9, t_4, 1]$ |
| {BAR, Iso} | {Iso} | $[p_{12}, p_{11}, t_4, 1]$ |
| {BAR, Gs, Iso, Gbg} | {Gs} | $[p_{26}, p_3, t_0, 1]$ |
| {BAR, Gs, Iso, Gbg} | {Gbg} | $[p_{26}, p_{17}, t_0, 1]$ |
| {BAR, Gs, Iso, Gbg} | {BAR, Iso} | $[p_{26}, p_{12}, t_{14}, 1], [p_{26}, p_{12}, t_0, 1]$ |
| {BAR, Gs, Iso, Gbg} | {Gs, Gbg} | $[p_{26}, p_2, t_{14}, 1]$ |
| {BAR, Gs, Iso, Gbg} | {BAR, Gs, Gbg} | $[p_{26}, p_{10}, t_{26}, 1]$ |
| {BAR, Gs, Iso, Gbg} | {Iso} | $[p_{26}, p_{11}, t_{26}, 1]$ |
| {BAR, Gs, Gbg} | {BAR, Gs, Iso, Gbg} | $[p_{10}, p_{26}, t_{25}, 2]$ |
| {BAR, Gs, Gbg} | {BAR} | $[p_{10}, p_9, t_{32}, 2]$ |
| {BAR, Gs, Gbg} | {Gs, Gbg} | $[p_{10}, p_2, t_{32}, 2]$ |
| {BAR} | {BAR, Gs, Gbg} | $[p_9, p_{10}, t_{31}, 2]$ |
| {BAR} | {BAR, Iso} | $[p_9, p_{12}, t_3, 2]$ |
| {Gs, Gbg} | {BAR, Gs, Iso, Gbg} | $[p_2, p_{26}, t_{13}, 2]$ |
| {Gs, Gbg} | {BAR, Gs, Gbg} | $[p_2, p_{10}, t_{31}, 2]$ |
| {Gs} | {AC, Gs} | $[p_3, p_0, t_{17}, 2]$ |
| {Gbg} | {Gs, Gbg} | $[p_{17}, p_2, t_{12}, 2]$ |
| {Gbg} | {GRK, Gbg} | $[p_{17}, p_4, t_6, 2]$ |
| {AC, Gs} | {AC} | $[p_0, p_{19}, t_{18}, 3]$ |
| {Gs} | {Gs, Gbg} | $[p_1, p_2, t_{12}, 3]$ |
| {AC, Gs} | {Gs} | $[p_0, p_3, t_{18}, 3]$ |
| {AC, Gs} | {cAMP} | $[p_0, p_{28}, t_{10}, 3]$ |
| {GRK, Gbg} | {BAR, Iso} | $[p_4, p_5, t_{30}, 3]$ |
| {GRK, Gbg} | {GRK} | $[p_4, p_{21}, t_7, 3]$ |
| {GRK, Gbg} | {Gbg} | $[p_4, p_{17}, t_7, 3]$ |
| {AC} | {AC, Gs} | $[p_{19}, p_0, t_{17}, 4]$ |
| {AC} | {cAMP} | $[p_{19}, p_{28}, t_{19}, 4]$ |
| {cAMP} | {cAMP, PKA} | $[p_{28}, p_{34}, t_{40}, 4], [p_{28}, p_{31}, t_{34}, 4], [p_{28}, p_{32}, t_{36}, 4], [p_{28}, p_3, t_{38}, 4]$ |
| {GRK} | {BAR, Iso} | $[p_{21}, p_5, t_{20}, 4]$ |
| {GRK} | {GRK, Gbg} | $[p_{21}, p_4, t_6, 4]$ |
| {cAMP, PKA} | {PKA} | $[p_{34}, p_{35}, t_{41}, 5]$ |
| {cAMP, PKA} | {PDE} | $[p_{31}, p_{30}, t_{21}, 5]$ |
| {cAMP, PKA} | {PTP} | $[p_{31}, p_{25}, t_{16}, 5]$ |
| {cAMP, PKA} | {BRaf} | $[p_{31}, p_{16}, t_9, 5]$ |
| {cAMP, PKA} | {cAMP} | $[p_{34}, p_{28}, t_{41}, 5], [p_{32}, p_{28}, t_{37}, 5], [p_3, p_{28}, t_{39}, 5], [p_{31}, p_{28}, t_{35}, 5]$ |
| {BRaf} | {MEK} | $[p_{16}, p_{15}, t_{22}, 6]$ |
| {PKA} | {cAMP, PKA} | $[p_{35}, p_{34}, t_{40}, 6]$ |
| {PDE} | {cAMP} | $[p_{30}, p_{20}, t_1, 6], [p_6, p_{20}, t_5, 7]$ |
| {PTP} | {MAPK} | $[p_{25}, p_{23}, t_{23}, 6], [p_{24}, p_{23}, t_{24}, 7]$ |
| {MEK} | {MAPK} | $[p_{15}, p_{13}, t_8, 7]$ |

**Figure S1:**
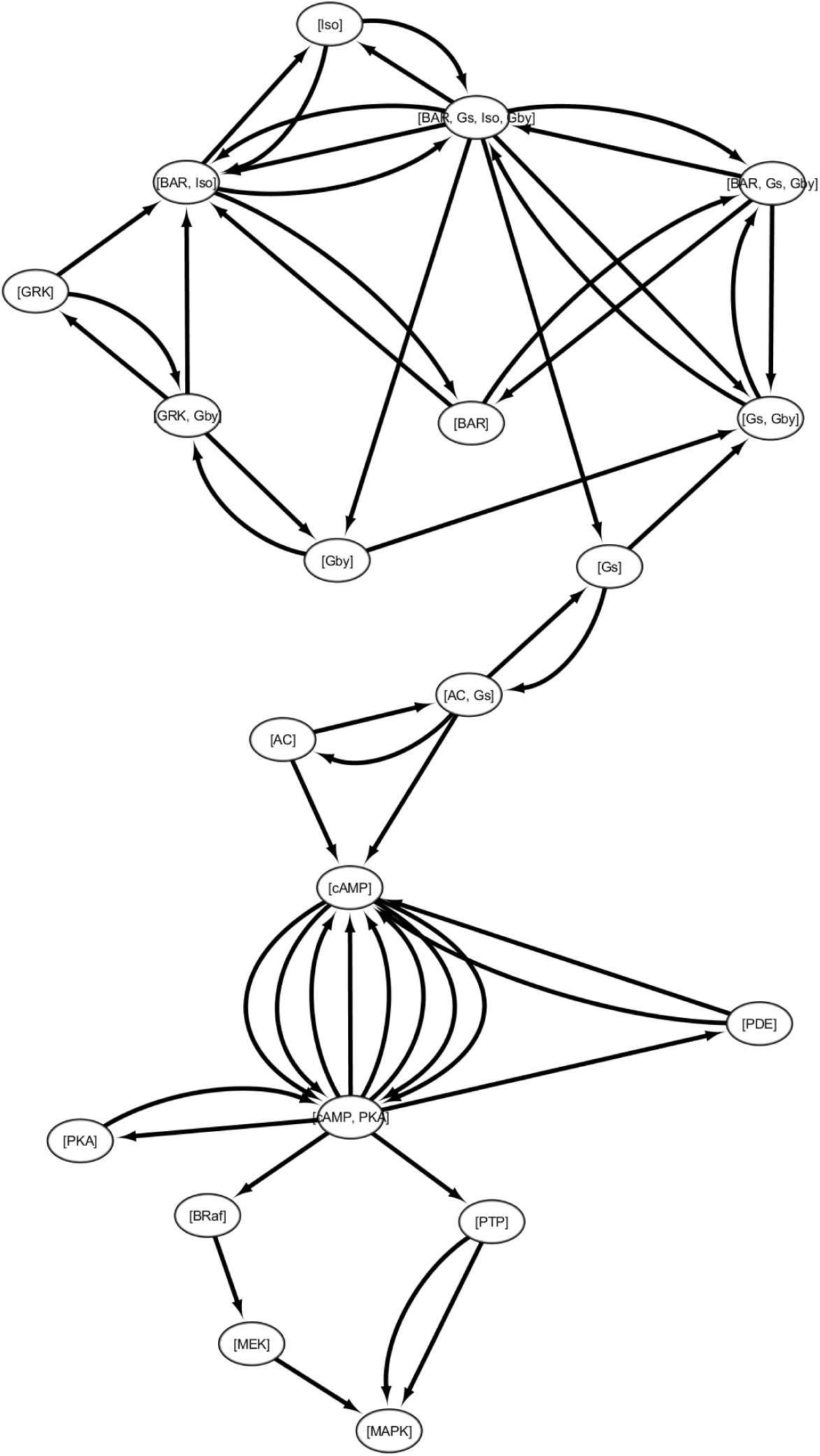
The interaction graph of the Petri net *N_GPCR_* as it is initially created by the first rule of the second phase of the algorithm with its vertices and their associated p-invariants. Table S2 lists the quadruplets associated with the edges of the graph.

**Figure S2:**
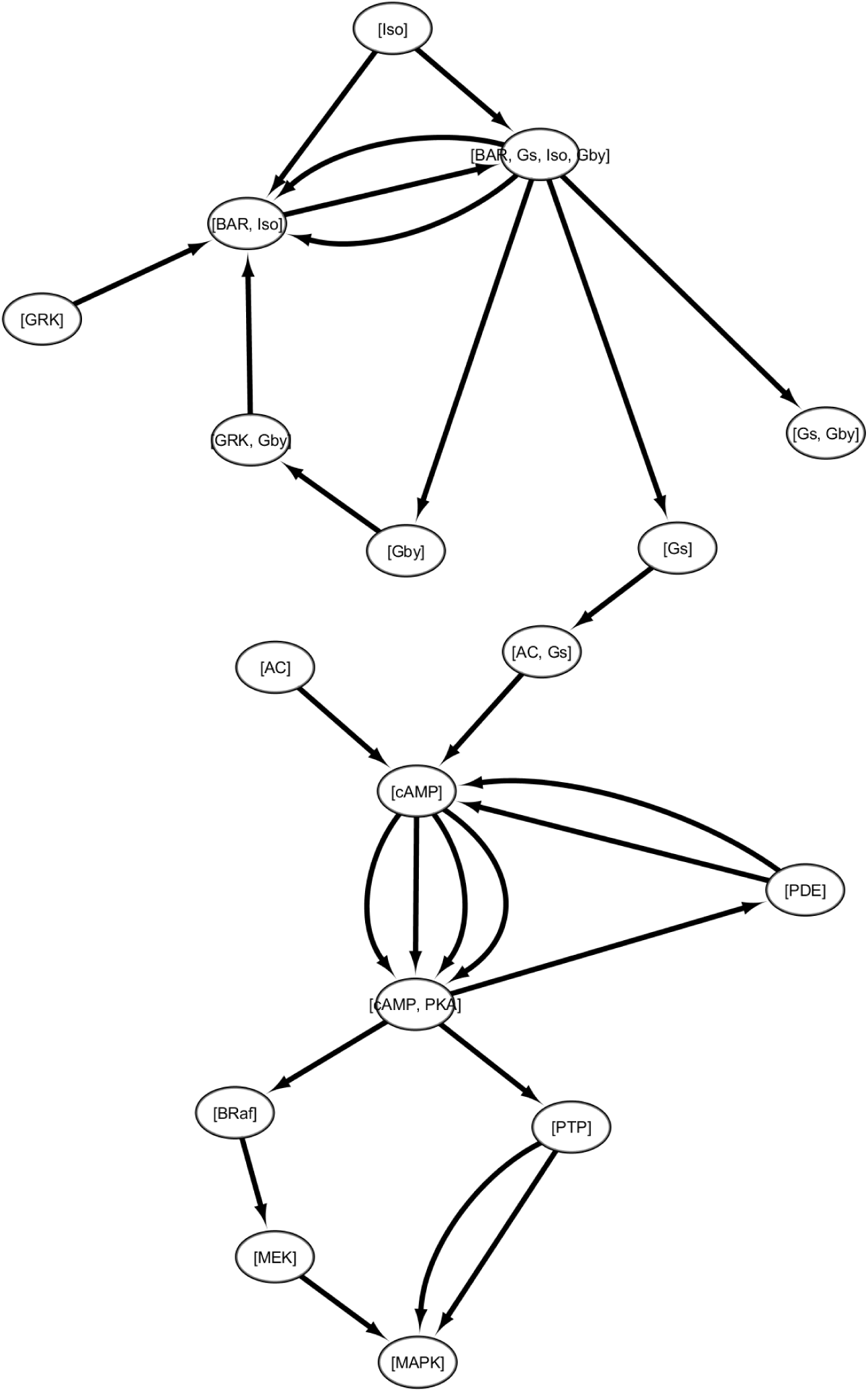
The interaction graph of the Petri net *N_GPCR_* after some edges are removed by the application of the signaling segments rule.

**Figure S3:**
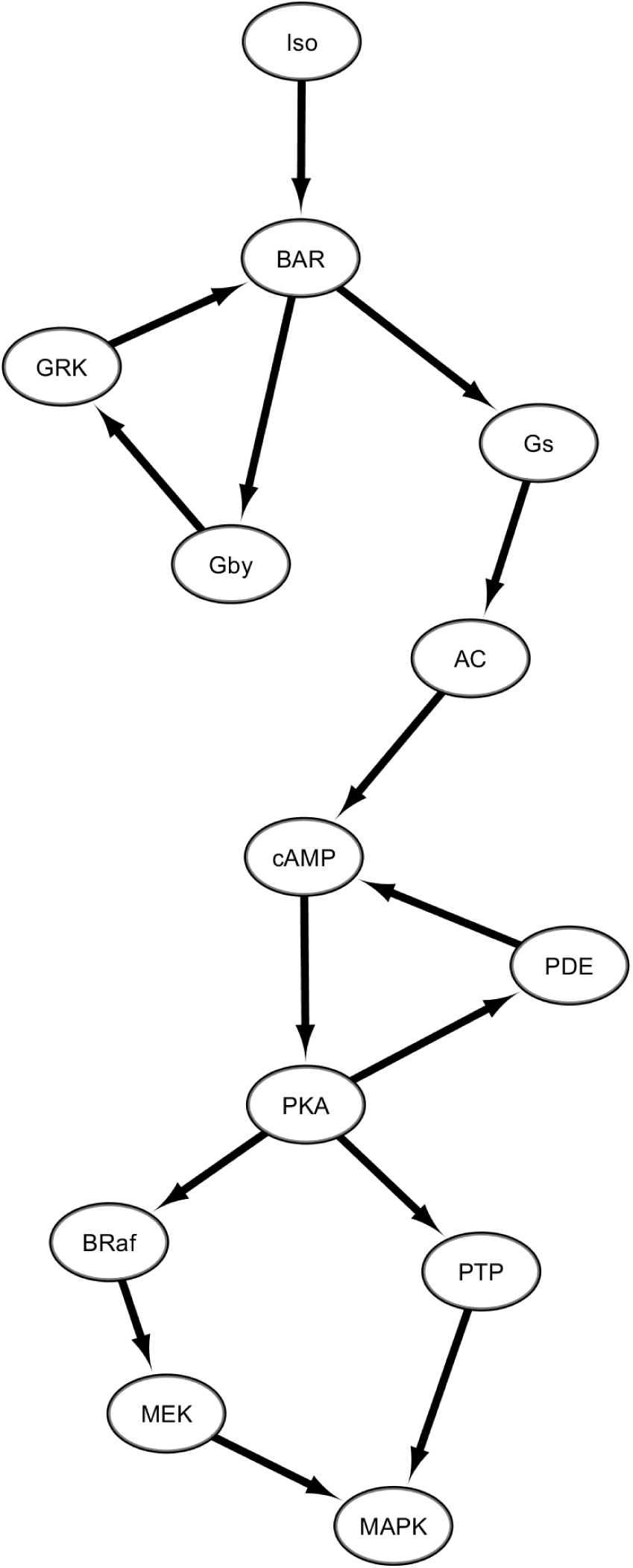
The unnanotated influence graph of the Petri net *N_GPCR_* after the execution of the third rule of the second phase of the algorithm.

